# Beyond magnetosomes: ubiquitous and diverse intracellular inclusions expand the role of magnetotactic bacteria in biogeochemical cycling

**DOI:** 10.64898/2025.12.17.695019

**Authors:** Qingxian Su, Bing Song, Alejandro Palomo González, Irini Angelidaki, How Yong Ng, Barth F. Smets, Kasper Reitzel, Bo Thamdrup, Marlene Mark Jensen

## Abstract

Magnetotactic bacteria (MTB) are capable of accumulating and storing intercellular pools containing phosphorus, sulfur, nitrogen and carbon. Yet, the metabolic pathways connected with their intracellular storage of polyphosphate, elemental sulfur, nitrate, and calcium carbonate, and the environmental influence on MTB inclusions and their quantitative contribution to sedimentary chemical budgets remain underexplored. Using a combination of chemical characterization, ultrastructural and compositional analyses, and phylogenetic and genomic insights into microbial assemblages, we investigated the influence of geochemical parameters on the diversity, ecophysiology, and distribution of MTB in freshwater and brackish sediments. Special focus is given to intracellular inclusions and their metabolic pathways to uncover functional traits and ecological roles in elemental cycling. Of the 118 examined MTB cells, polyphosphate granules (75%) and nitrate-containing vacuoles (67%) were the most common inclusions followed by sulfur globules (25%) and calcium carbonate granules (8%). The intracellular phosphorus, nitrogen, and sulfur stored in MTB cells were conservatively estimated to account for 0.60%, 0.91%, and 0.23% of the total sedimentary P, N, and S in the investigated freshwater sediments, respectively. The ubiquitous nature and important ecological role of MTB can be explained by their ability to sequester chemical elements into intracellular inclusions, giving them a metabolic advantage in dynamic chemically stratified environments. The extraordinary phylogenetic diversity of MTB, coupled with their capability to hyperaccumulate and store a wide range of elements and compounds, represents a significant resource for biotechnology innovation.

## Introduction

Magnetotactic bacteria (MTB) are a genetically and morphologically diverse group of microorganisms unified by their ability to biomineralize magnetosomes. Magnetosomes are magnetic nanocrystals in the form of magnetite (Fe_3_O_4_) and/or greigite (Fe_3_S_4_) enveloped by a lipid bilayer membrane ^1,2^. Using a behavior known as magnetotaxis, the magnetosomes enable MTB to align and navigate along Earth’s magnetic field, providing path through vertically stratified chemical gradients to find zones with favorable redox conditions ^3^. While MTB are ubiquitous in sediments and water columns of freshwater, brackish, marine, and hypersaline habitats, they have proven difficult to grow in the laboratory and their wide taxonomic diversity in natural samples is not represented as isolates ^3–6^. Most cultured and uncultured MTB are affiliated with Alphaproteobacteria, Gammaproteobacteria, Desulfobacterota, Nitrospirota, Planctomycetes, and the candidate phyla Omnitrophica and Latescibacteria ^6–8^. MTB are usually detected below and near the oxic-anoxic transition zone ^3,9,10^. The type of magnetic nanocrystal might influence the location of MTB or vice versa, as the formation of Fe_3_S_4_ crystals is favored under sulfidic conditions, while Fe_3_O_4_ formation requires oxygen (O_2_) ^10^.

MTB have long been considered important in iron (Fe) cycle, as they take up dissolved Fe from the environment during biomineralization, contributing to the burial of Fe in the sediments ^11,12^. While the diversity and mechanism of biomineralization of magnetosomes have been extensively studied, much less is known about the metabolic versatility that enables MTB to inhabit distinct biogeochemical niches and thereby contribute to different biogeochemical cycles. Genomic-based studies have provided information on the metabolic potentials of uncultured MTB, especially the ones affiliated with the phylum Nitrospirota ^13–16^. For instance, genomic analyses of *Ca.* Magnetobacterium casensis and *Ca.* Magnetobacterium bavaricum indicated an autotrophic lifestyle together with capabilities for denitrification, sulfur oxidation, and/or sulfate (SO_4_^2−^) reduction ^13,14,16^. Furthermore, this metabolic versatility enables them to actively concentrate chemical elements of sulfur (S), phosphorus (P), and nitrogen (N) in intracellular inclusions, identified as S globules, polyphosphate (polyP) granules, and vacuoles possibly with nitrate (NO_3_^−^) ^6,13,15,17–21^. Inclusions with polyP have been frequently observed among freshwater and marine magnetotactic cocci ^17,20,22–24^. For instance, by filling most of their cytoplasm with polyP granules, magnetotactic cocci in the water column of Black Sea was conservatively estimated to sequester over 6 fmol of P per cell – a storage capacity over 100 times greater than that of ordinary-sized bacteria like *Sulfurimonas gotlandica* ^20^. This remarkable storage capacity, coupled with their motility, enables MTB to function as active “bacterial phosphate shuttles”, potentially taking up phosphate (PO_4_^3^⁻) near the oxycline and releasing it at the lower boundary of the oxygen-deficient zone. Sulfur globules are found in MTB affiliated with the Eta-, Alpha-, and Gamma-proteobacteria classes, the Nitrospirota phylum, and the candidate phylum Omnitrophica ^14,25,26^. For example, both S globules and NO_3_^−^-filled vacuoles were observed in *Ca.* Magnetobacterium casensis cells (phylum Nitrospirota), which were speculated to shuttle NO_3_^−^ downward in the sediment for reduction and sulfur upward for oxidation ^13^. Apart from Fe, polyP, S, and NO_3_^−^, some MTB also harbor granules containing polyhydroxyalkanoates (PHA) (e.g., *Magnetospirillum* genus) and calcium carbonate (CaCO_3_) (e.g., Azospirilllaceae family) ^27–29^. In general, the physiological function of intracellular inclusions in MTB remains rather unclear. In phylogenetic unrelated bacteria, polyP- and PHA-inclusions, S globules, and NO_3_^−^-storing vacuoles serve as storage depots for elements and/or energy ^30^. For instance, in enhanced biological phosphate removal (EBPR) of wastewater treatment trains, polyphosphate-accumulating organisms (PAOs) store polyP under oxic conditions by oxidizing PHA as an energy source, and subsequently degrade polyP under anoxic conditions to generate energy for rebuilding their intracellular PHA pool ^31^. Sulfur bacteria, such as *Beggiatoa*, *Thioploca*, and *Thiomargarita*, can accumulate reduced sulfur in S globules during aerobic sulfide oxidation to sustain anaerobic metabolism and growth ^32–34^. In the marine representatives of *Beggiatoa*, NO_3_^−^ is stored in vacuoles, which is used as an electron acceptor for anaerobic sulfide oxidation in deep sediment layers ^35^. Thus, in both sulfur bacteria and PAOs, it is the availability and distribution of O_2_, sulfur compounds, and NO_3_^−^that seem to drive the synthesis and degradation of inclusions in these bacteria ^36,37^. As a group of prokaryotes capable of moving up-down within the oxic-anoxic transition zone with the help of magnetotaxis, MTB may sequester chemical elements (i.e., Fe, N, S, and P) into cells in response to the stratification of specific geochemical parameters. However, the link between geochemical parameters and intracellular inclusions in MTB remains to be elucidated.

MTB act as important engines that drive global Fe cycling, yet their impact on the other biogeochemical cycles is less understood. A comprehensive understanding of MTB metabolism and their adaptation to environmental conditions is required to assess their ecological roles, particularly in the biogeochemical cycles, as well as developing new strategies to exploit MTB for beneficial biotechnology application. This paper will explore how geochemical conditions shape the diversity, ecophysiology, and distribution of MTB in brackish and freshwater sediments. We give attention to the intracellular inclusions (including polyP granules, NO_3_^−^ vacuoles, S globules, and CaCO_3_ inclusions) and the associated metabolic pathways with the aim of exploring functional traits of MTB and elucidating their ecological role in P, N, and S cycling.

## Results and Discussion

### Distinct features in physical-chemical characteristics and microbial landscapes

The sediments at three freshwater sites (Östra Silen, Furesø, and Lake Ormstrup) and two brackish sites (Flensborg Fjord and Faxe Bugt) were characterized by distinct physical and chemical features (Table S1–S2, Fig. S2). Organic matter mineralization rates measured in Flensborg Fjord (32 ± 4.3 mmol/m^2^/d) and Faxe Bugt sediments (12 ± 1.8 mmol/m^2^/d) were within the range of previous measurements performed in permeable sediments (Table S2) ^38^. These rates depend primarily on the quantity and quality of organic matter ^39^. However, unlike cohesive sediments, where diffusive transport limits O_2_ penetration to a few millimeters, the advective transport in sandy sediments in Flensborg Fjord and Faxe Bugt, where sampling sites were located within the subtidal zone, led to extended oxic and suboxic zones. A possible effect of advective porewater transport was indicated by the lower concentrations of hydrogen sulfide (H_2_S) and PO_4_^3^⁻ at 3–4 cm depth in Flensborg Fjord (Fig. S2). There was no obvious build-up of H_2_S in the porewater at any of the sites, except in Flensborg Fjord, indicating the highly reduced condition in the sediments. The 2–4 times higher mineralization rates in Lake Ormstrup sediments compared to Furesø and Östra Silen reflected the eutrophic state of the lake and the high amount of labile organic matter (Table S2). Together with organic matter, the high concentrations of ortho-PO_4_^3^⁻ in Lake Ormstrup sediments were due to a large accumulated P pool originating from previous discharge of wastewater, feeding of ducks for hunting purposes, and agriculture ^40^. The highly reduced conditions in Lake Ormstrup, especially at the deep site, where the above water column was stratified during our sampling, were confirmed by the anoxic bottom water and high concentrations of ammonium (NH_4_^+^) (7–8 times higher), Fe^2+^ (7–24 times higher), Fe(II) (6–18 times higher), acid volatile sulfide (AVS) (10–116 times higher), and chromium reducible sulfur (CRS) (8–31 times higher), compared to the other freshwater sediments (Table S2, Fig. S2). The notably high AVS and CRS concentrations in Ormstrup sediments compared with the other freshwater sites were broadly consistent with previous studies, where a positive relationship between pools of AVS and CRS and increasing eutrophication was observed ^41–43^. Concentrations of PO_4_^3^⁻ detected in Östra Silen (0.025 mmol/m^2^) were significantly lower than in Lake Ormstrup and Furesø (0.24–4.3 mmol/m^2^). Furthermore, the sediment in Östra Silen was highly enriched in FeOOH (132 mmol/m^2^), which derived from the input by streams from leached acidic pine forest soils and peat lands ^44,45^. Elevated levels of FeOOH caused a decrease in porewater PO_4_^3^⁻ concentrations and flux out of the sediment, and ultimately resulted in an oligotrophic state of Östra Silen. The sediment in Östra Silen was Ca poor (Table S2), which often characterizes dystrophic lakes due to being in a Ca-poor area or high content of humic substances ^46^.

Different environmental factors influence microbial community assembly. Brackish sediments were enriched in microbial clades associated with S cycling (e.g., Desulfobacterales and Chromatiales), while freshwater sites featured methane-oxidizing groups (e.g., Methylococcales and Methylomirabilales), consistent with published studies on sediment microbiomes (Fig. S4) ^47,48^. Notably, sediments in Östra Silen exhibited the highest microbial diversity, likely due to the greater niche differentiation under nutrient-poor (oligotrophic) state^49^. Furthermore, microbial composition changed significantly with sediment depth at all sites, where the dominant microbes shifted from aerobic methane oxidizers (e.g., Methylococcales) and photoheterotrophs (e.g., Rhodobacterales) in surface sediments to anaerobic fermenters (e.g., Clostridiales and Syntrophobacterales), anaerobic methane oxidizers (Methylomirabilales), and sulfate reducers (e.g., Desulfobacterales and Desulfarculales) in the deeper sediment layers (Fig. S4). The variation of microbial composition with sediment depth reflects the influence of chemical conditions, nutrient availability, and redox potential, which together shapes the community structure and diversity. While the microbial composition of Östra Silen and Furesø sediments showed a clear gradual transition from aerobic to anaerobic microbes with depth, this shift was less variable in Lake Ormstrup, likely linked to the anoxia in bottom water and strongly reduced conditions in the sediments.

The maximum likelihood phylogenetic tree based on 16S rRNA gene sequences of cells purified using a magnetic separation technique at each site resulted in 36 different Amplicon Sequence Variants (ASVs) affiliated with the *Magnetococcus* genus of Alphaproteobacteria (Fig. 1A). There was a clear clustering of brackish and freshwater *Magnetococcus* taxa, with 24 ASVs in brackish sediments and 12 ASVs in freshwater sediments. Some magnetotactic cocci found in the study may represent novel species due to the < 97% sequence similarity with the nearest known relative. While MTB affiliated with *Magnetococcus* were identified at all sites, taxa affiliated with *Magnetospirillum* occurred predominantly in freshwater sites (Fig. 1A). The majority of the identified *Magnetospirillum* MTB (e.g., 16s_108, 16s_109, 16s_24857, 16s_8465, and 16s_10249) was closely related (a similarity threshold >97%) to *Magnetospirillum sp.* RSS-1, *Magnetospirillum magnetotacticum*, *Magnetospirillum sp.* ME-1, *Magnetospirillum marisnigri*, *Magnetospirillum sp.* XM-1, and *Magnetospirillum gryphiswaldense*, all of which have been previously isolated from freshwater environments ^50–53^. The relative abundance of *Magnetococcus* and *Magnetospirillum* in the sediments was 0.002–0.012% and 0.06–0.1% of the microbial community, respectively, comparable to typical reported MTB abundance (< 0.01%) (Fig. 1B) ^4,54,55^. This agrees with many biogeographic studies of MTB, where either *Magnetococcus* or *Magnetospirillum* was identified as the most abundant genus ^3,7,56–59^. A single ASV found in the sediment of Östra Silen was closely related to *Ca.* Magnetominusculus xianensis in the phylum Nitrospirota (Fig. 1A) ^60^. ASVs affiliated with multicellular magnetotactic prokaryotes (MMP) were solely found at the brackish sites, consistent with their known prevalence in sulfur-rich brackish to marine environments ^21,61–64^. The multicellular MTB detected at Flensborg Fjord and Faxe Bugt shared a 16S rRNA similarity of > 97% with *Ca.* Magnetomorum sp. HK-1 and *Ca.* Magnetomorum sp. 1–7 within the Desulfobacterota, which are uncultivated spherical MMP producing both magnetite and greigite magnetosomes ^65,66^.

**Fig. 1.**
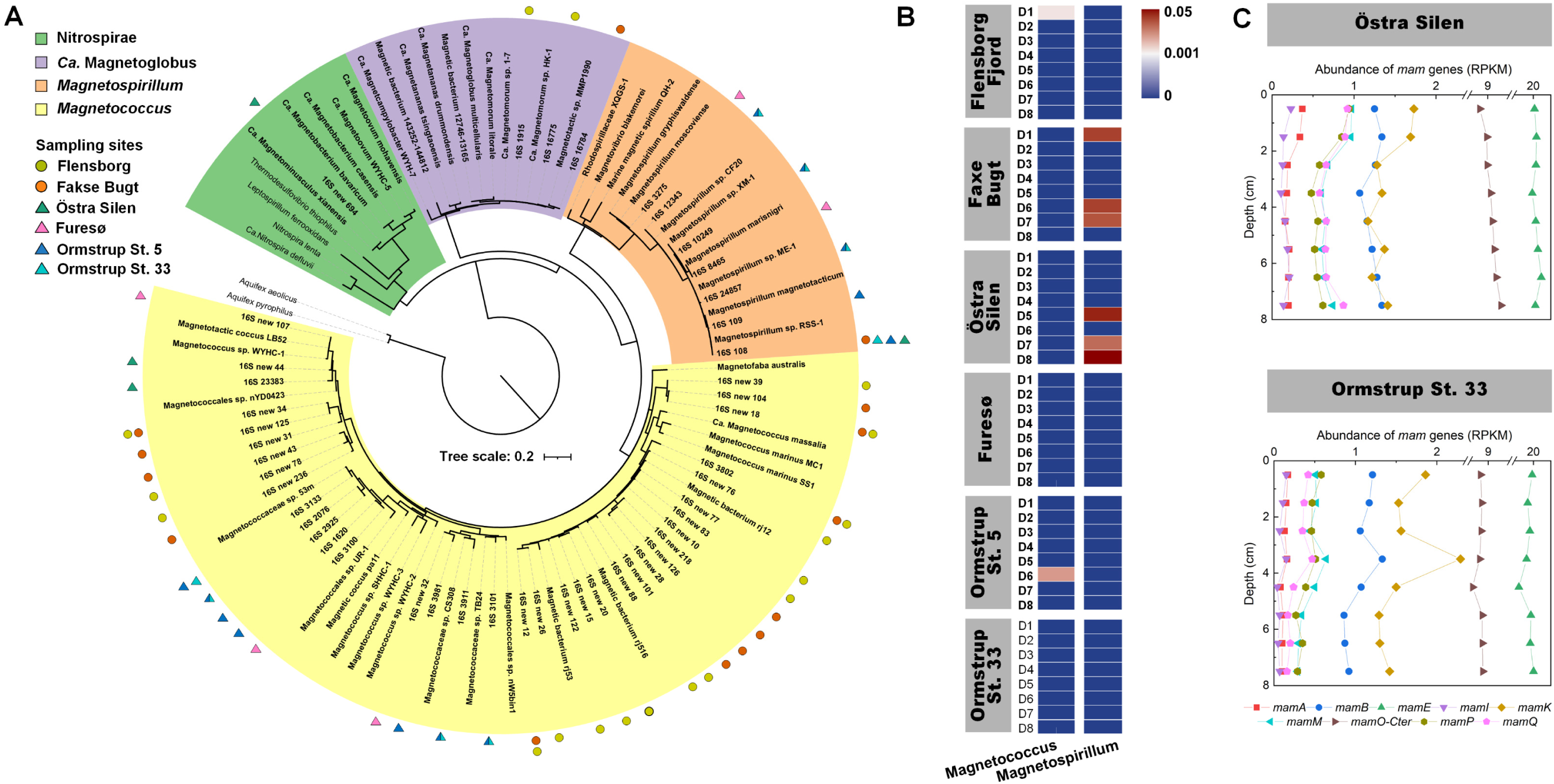
(A) Most-likelihood phylogeny of MTB. The phylogenetic tree was constructed based on 16S rRNA gene sequences of magnetically collected MTB cells. Mean Scale bars represent 0.2 μm. (B) Depth distribution of MTB. The DNA extracted from sediment samples at each depth interval was used for 16S rRNA gene sequence analysis. *Ca.* Magnetoglobus and Nitrospirae were not detected. (C) Depth distribution of the relative abundance of *mam* genes in the sediments of Östra Silen and Lake Ormstrup St. 33. The quantification of *mam* genes was determined based on metagenomic analysis. Abundance was given in reads per kilobase sequence per million mapped reads (RPKM).

We were able to obtain eight medium- to high-quality draft genomes for members of several abundant MTB taxa described above by 16S amplicon sequencing (Fig. S5, Table S5). Two draft genomes of MTB (LS_bin4 and LS_bin5) recovered magnetically from the sediments in Östra Silen and Furesø were putatively affiliated with the genus *Magnetospirillum* within the class Alphaproteobacteria, consistent with the 16S rRNA gene sequencing results (Fig. 1A, Table S5). The assembled draft genomes, BS_bin2 retrieved from Flensborg Fjord and Faxe Bugt, and LS_bin6 retrieved from Furesø, were assigned to the genus *Magnetomorum* and *Desulfamplus*, respectively, within the class *Desulfobacteria*. The remaining genomes of marine and primarily freshwater MTB (BS_bin1, LS_bin1, LS_bin2, and LS_bin3) found in Flensborg Fjord, Faxe Bugt, and Östra Silen were assigned to the class of Magnetococcia.

### Vertical distribution of MTB across chemical gradients

MTB are indigenous in sediments or chemically stratified water columns, where they occur predominantly at the oxic-anoxic interface, the anoxic regions, or both ^3,9,10^. In this study, we observed that MTB affiliated with *Magnetococcus* and *Magnetospirillum* were present both in the surface sediment and below 4 cm (Fig. 1B). Consistently, the abundance of magnetotactic cocci has been reported to vary by several orders of magnitude over a depth of 30 cm in salt marsh sediment, with maximum cell counts at the oxygen-sulfide interface ^9^. Moreover, cocci and spirilla MTB have been reported to exist in both oxic and anoxic zones of different freshwater sediments, with the largest proportion (63–98%) detected within the anoxic zone ^10^. As most of the cultivated MTB, especially *Magnetospirillum* strains, are known to behave as typical microaerophiles ^67^, their dominance in the deeper parts of the sediment remains puzzling (Fig. 1B). As gradient organisms, most MTB derive energy for growth from reductants and oxidants at a chemical interface, requiring them to appear at particular locations relative to this interface ^4^. The availability of inorganic electron donors and acceptors is considered an important factor determining the vertical distribution of MTB ^10^. It is therefore speculated that the occurrence of uncultivated *Magnetospirillum* in deeper layers of sediments could be due to its utilization of different, as yet unidentified electron acceptors.

Magnetosome membrane (*mam*) genes necessary for the biosynthesis and organization of magnetosomes are unique to MTB. The nine genes, *mamA*, *В*, *M*, *K*, *P*, *Q*, *E*, *O*, and *I*, annotated in the acquired draft genomes, have been identified as conserved across the majority of MTB genomes (Fig. 1C) ^25,68,69^. We observed significant correlations of the genes *mamM*, *mamO*, *mamP*, and *mamQ*, which are responsible for nucleation and growth of magnetite crystal, with concentrations of TOC, TN, TP, Fe(II), SO_4_^2−^, and H_2_S (*p* < 0.05) (Fig. S8). Together with the significant correlation between *mam* genes and concentrations of CRS and AVS (*p* < 0.05) in the depth profiles of Östra Silen and Lake Ormstrup, these associations with sulfur compounds indicate a strong dependence of MTB on sulfur compounds in their metabolisms (Fig. S8). The ability to oxidize sulfur compounds, likely as electron donors, has been observed among MTB^10^. Our observations are consistent with previous biogeographic studies, where the relative abundance of MTB in brackish sediments correlated with the concentrations of organic matter, elemental sulfur (S^0^), SO_4_^2−^, H_2_S, NH_4_^+^, NO_3_^−^, and PO_4_^3^⁻ ^9,55,70,71^. The distribution of MTB across multiple chemical gradients, as exemplified by the *Magnetospirillum* in Östra Silen and Faxe Bugt, indicates that the distribution of MTB is not controlled by a single factor but rather a ‘‘trade-off’’ between the availability of nutrients, electron donors, and electron acceptors ^10^.

### Comparative diversity of cell morphotypes and intracellular inclusions across habitats

Molecular typing methods, such as the FISH-SEM approach, are required to link MTB phylogeny with morphology ^72^. However, the unique ultrastructure of certain MTB allows for informed speculations about their taxonomy ^58^. Our scanning transmission electron microscopy (STEM) images revealed a diverse range of cell morphotypes, including both habitat-specific and cosmopolitan types (Fig. 2). Distinct environmental conditions selected for specific populations of MTB (Fig. 1A). This is especially true for *Desulfobacteria*-affiliated MMP, which with their distinct cell morphology are typically observed only in saline aquatic habitats ^21,61–64^. Based on light microscopy observations, spherical mulberry-like MMP (sMMP) were visually abundant in Flensborg Fjord, each composed of 27–53 cells (e.g., Fig. 2 A1-A3). All MMP have been reported to mineralize magnetosomes consisting of Fe_3_O_4_ and/or Fe_3_S_4_ crystals ^61,65,70^. MMP found in Flensborg Fjord were affiliated with the genus *Ca.* Magnetomorum within the class *Desulfobacteria* ^65,70,73^. The draft genome BS_bin2, affiliated with *Ca.* Magnetomorum was likely the MMP visualized in images A1-A3 of Fig. 2. Furthermore, genes associated with dissimilatory sulfate reduction were identified in BS_bin2, including sulfate adenylyltransferase (*sat*), adenylyl-sulfate reductase (*apr*), and dissimilatory sulfite reductase (*dsr*) (Fig. 4). This substantiates the sulfate-reducing lifestyle of *Ca.* Magnetomorum ^74,75^, and supports the observed strong correlation of MTB distribution with sulfur compounds (Fig. S2, Fig. S8).

**Fig. 2.**
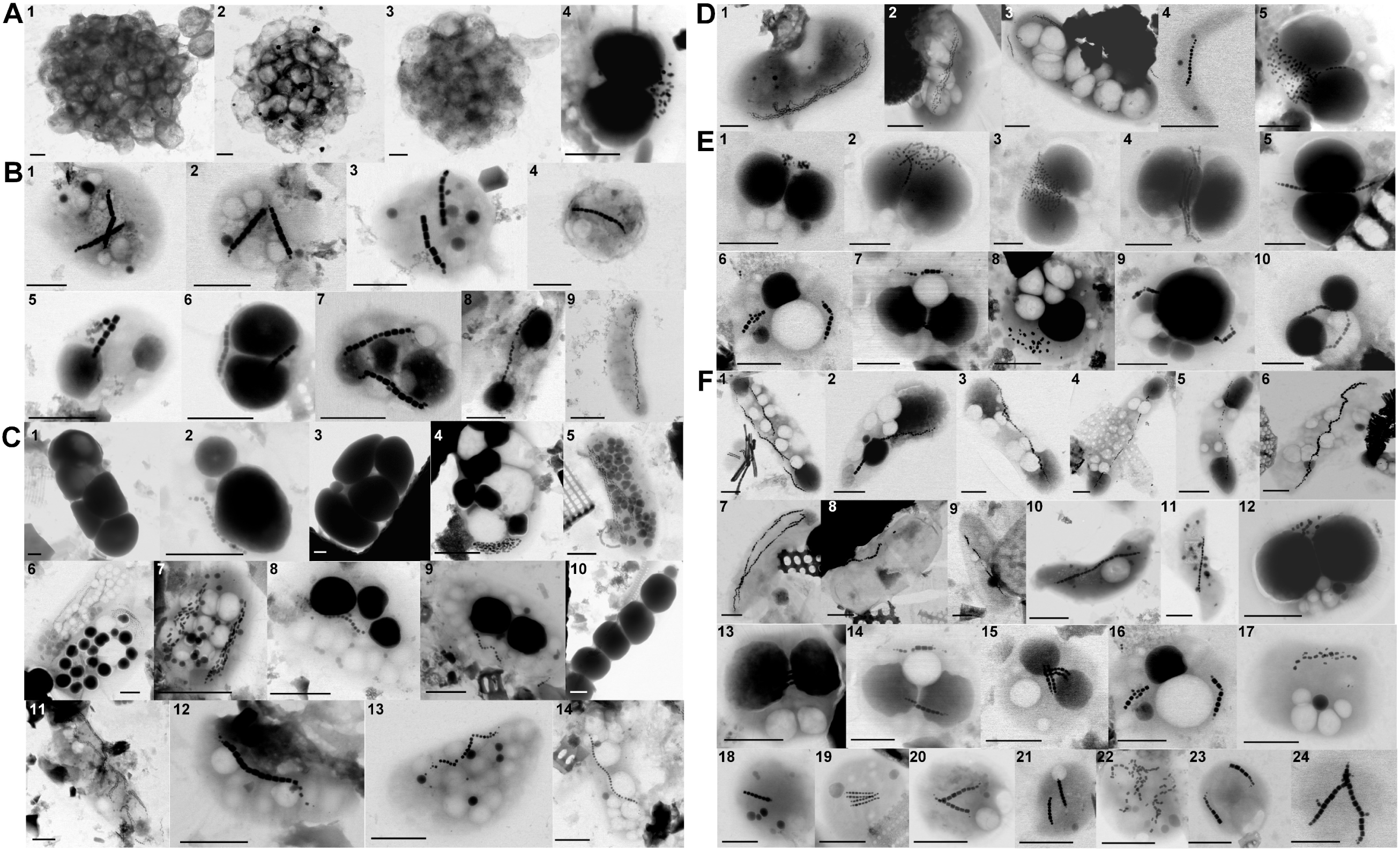
Examples of scanning transmission electron microscopy (STEM) images of magnetically enriched MTB from (A) Flensborg Fjord, (B) Faxe Bugt, (C) Östra Silen, (D) Furesø, (E) Lake Ormstrup St. 5, and (F) Lake Ormstrup St. 33. Scale bars represent 1 μm. The diameter and area of MTB cells and their intracellular inclusions are presented in Supplementary Information Table S3.

Magnetotactic cocci are cosmopolitans and often found to be the most frequent morphotype in both freshwater and marine habitats ^3,58^. In consistency with our results, magnetotactic cocci were highly abundant, comprising ∼76% of magnetically collected cells (Fig. 1A), and visually observed across all our sampling sites, with varying size of 1.3–9.9 μm^2^ (Fig. 2, Fig. S7). These coccoid-shaped cells were likely affiliated with *Magnetococcus* ^58,76^. The presence of at least ten morphologically distinct magnetotactic cocci in Lake Ormstrup is consistent with previous findings that different populations of magnetotactic cocci often dominate in muddy sediments with a relatively high content of organic material (Table S1–S2) ^4,56^. The magnetotactic spirilla found at Station 33 in Lake Ormstrup (Fig. 2 F10-F11) closely resembled MTB affiliated with the *Magnetospira* genus in Lake Pavin ^77^.

At least 10 morphologically distinct MTB were found in Östra Silen (Fig. 2). Multiple rods producing bullet-shaped magnetosomes, consisting of a few hundred bullet-shaped magnetic crystals, were only observed in Östra Silen (Fig. 2 C5-C6, C11). These cells, ranging between 1.5–2.0 μm in width and 4.6–6.9 μm in length, are morphologically similar to previously identified strains, *Ca.* Magnetobacterium casensis ^16^ and *Ca.* Magnetobacterium bavaricum ^26^, both affiliated with Nitrospirota. Additional special rods containing aligned/nested large refractive inclusions occupied the majority of the cytoplasm were only detected in Östra Silen (e.g., Fig. 2 C1-C3, C9-C10, Fig. S9). These morphotypes are comparable to the reported CaCO_3_-storing MTB affiliated with the family of Azospirilllaceae and CAIRSR01 of Gammaproteobacteria ^28,29^. In summary, environmental heterogeneity was found to explain the variation in diversity and distribution of MTB in different sampling sites, indicating that environmental conditions are crucial in structuring MTB community composition. Salinity and sulfide appear to be key factors shaping the distribution of MMP ^21,61,63^ (Fig. 2), while the low concentrations of Ca might select for CaCO_3_-storing MTB in Ostra Silen ^27–29^. In general, the high morphological diversity in Ostra Silen might be linked to the multiple stable chemical micro-gradients (Fig. 2, Fig. S2) ^4,55^. The widespread presence of magnetotactic cocci is intriguing. The *Magnetococcus* species appear to harbor larger and different types of inclusions compared to many other MTB ^15,17,18,20,78^. The ability to store and potentially metabolize multiple compounds like P, S, and N might give *Magnetococcus* species a competitive advantage in stratified nutrient-variable environments.

### CaCO3 granules

In the sediment of Ostra Silen, electron-dense inclusions in MTB were characterized by high intensities of Ca, C, and O, indicating that they were likely composed of CaCO_3_ (Fig. 2–3). Similar Ca-rich granules have been found in MTB affiliated to the Azospirilllaceae family within the Alphaproteobacteria, including a novel strain XQGS-1 isolated from the freshwater sediments of Xingqinggong Lake ^27^ and a magnetotactic slightly-curved rod (named CCP1) in the ferruginous Lake Pavin ^28^. Based on an ultrastructural characterization using TEM and synchrotron-based scanning transmission X-ray microscopy (STXM), the granules in MTB from Lake Pavin were composed of amorphous CaCO_3_ and contained within membrane-delimited vesicles ^28^. The STEM images revealed four different morphotypes of CaCO_3_ inclusions (n = 48): (1) rod-shaped cells with 2–4 large inclusions, corresponding to 70 ± 16% of the cytoplasmic (n = 20), (2) elongated cells (up to 10 μm in length) containing 4–5 aligned inclusions, (3) cells with large, irregular, nested inclusions filling the cytoplasm and overlapping magnetosomes, and (4) cells harboring 2–17 unaligned spherical granules, occupying 38 ± 0.10% of the cytoplasmic space (Fig. 2C, Fig. S9). Our 16S rRNA gene sequencing and metagenomic analyses did not cover the phylogeny and genomes of CaCO_3_-storing MTB in Östra Silen sediments. We found that CaCO_3_ inclusions in morphotypes 1–3 closely resembled those reported by Mangin and coworkers (2025) ^29^ (Comparison is shown in Fig. S9). Thus, with respect to phylogeny and metabolic pathways associated with CaCO_3_ formation, we refer to the results in Mangin et al. (2025) ^29^. The single-cell amplified genome sequencing of different morphotypes of CaCO_3_-storing MTB revealed that each morphotype was associated with a genomic group affiliated with divergent families within the Pseudomonadota: Morphotypes 1–2 and 3 were affiliated with the Azospirilllaceae family and the family CAIRSR01 of Gammaproteobacteria, respectively ^28,29^. Metabolic network modeling of CaCO_3_-storing MTB belonging to Gammaproteobacteria indicated the presence and potential utilization of the Calvin-Benson-Bassham (CBB) cycle for CO_2_ fixation and intracellular carbonate formation, associated with the oxidation of reduced sulfur compounds (mainly S^2−^ and thiosulfate (S_2_O_3_^2−^)) ^29,79,80^. In contrast, CaCO_3_-storing MTB belonging to Azospirillaceae family predominantly utilize the reductive citrate (rTCA) cycle for CO_2_ fixation, and has the potential to perform NO_3_^−^ and nitrite (NO_2_^−^) reduction ^29^. The energy for CaCO_3_ biomineralization might be gained from sulfur oxidation by using O_2_ or NO_3_^−^ as potential electron acceptors ^81,82^. However, the metabolic pathways underlying the synthesis and degradation of CaCO_3_ by MTB remain insufficiently characterized and call for further investigation.

**Fig. 3.**
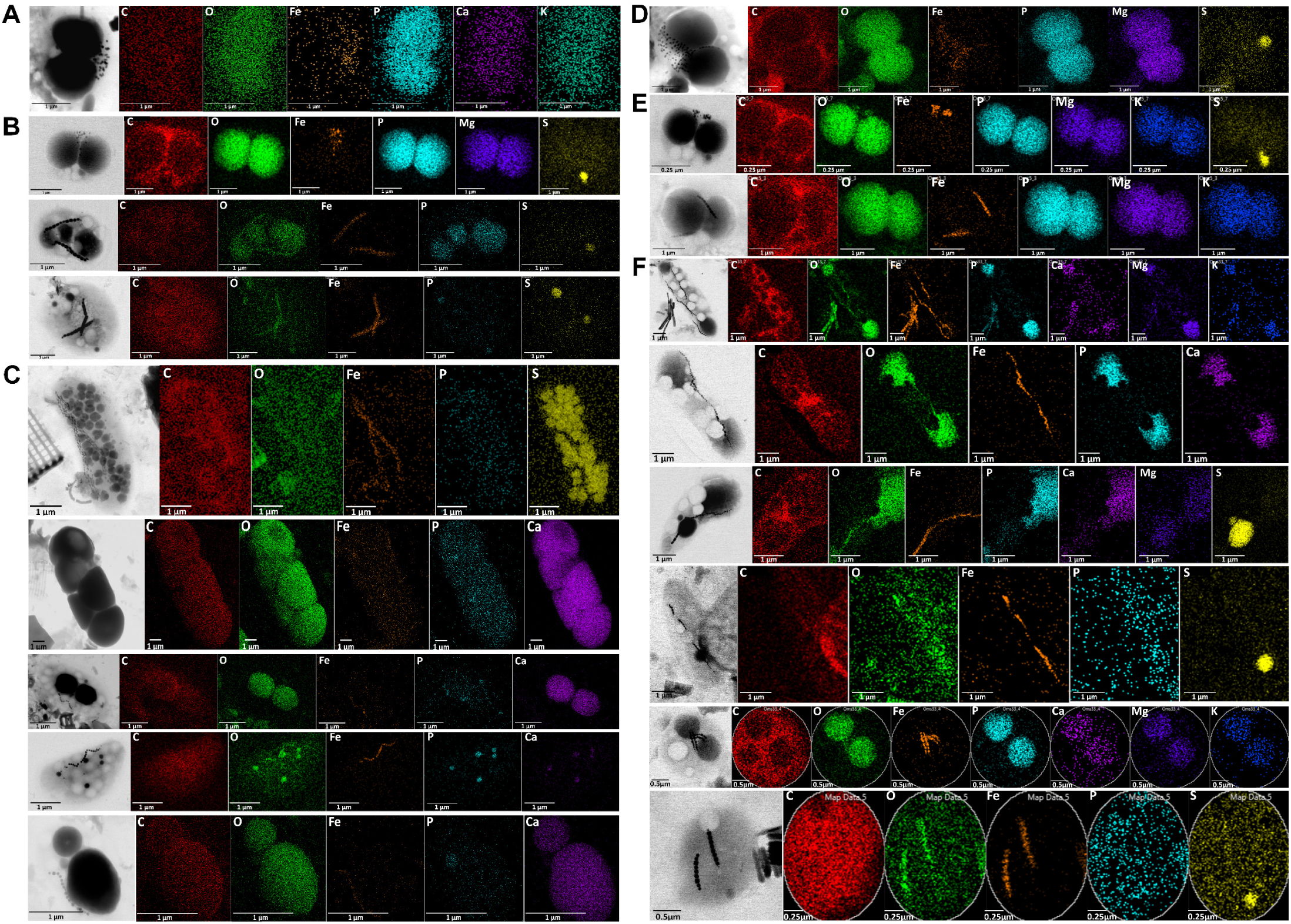
Examples of STEM-EDX elemental mapping of carbon (C), oxygen (O), iron (Fe), phosphorus (P), sulfur (S), calcium (Ca), Magnesium (mg), and potassium (K) in magnetically enriched MTB cells from (A) Flensborg Fjord, (B) Faxe Bugt, (C) Östra Silen, (D) Furesø, (E) Lake Ormstrup St. 5, and (F) Lake Ormstrup St. 33.

The potential functions of intracellular CaCO_3_ granules may be involved in (i) the storage of inorganic carbon that may serve as an electron acceptor in carbonate respiration, (ii) a buoyancy-regulating mechanism, or (iii) the buffering of intracellular pH ^81,82^. We exclusively found CaCO_3_-rich granules in MTB from Östra Silen (Fig. 2–3, Fig. 5A). The lake was characterized by high concentrations of solid Fe and low concentrations of dissolved Ca^2+^, i.e., 12–42 times lower Ca^2+^ concentrations than other sampling sites (Fig. S2, Table S2). Consistently, MTB in the ferruginous Lake Pavin were observed to store CaCO_3_ intracellularly in sediment layers where porewaters were undersaturated with CaCO_3_ phases ^28,29^. As these environmental conditions are thermodynamically unfavorable for mineral precipitation, forming CaCO_3_ granules in MTB is likely an active process, and is energetically costly to maintain an intracellular environment supersaturated with CaCO_3_ ^29,83^. Although the two lakes share some common environmental characteristics – both being oligotrophic, Fe-rich, and Ca^2+^-poor – the environmental determinants underlying the preferential proliferation of CaCO_3_-storing MTB in these specific habitats remain largely undefined. Addressing this research gap necessitates a comprehensive investigation into the unique geochemical conditions of these environments, the specific metabolisms of the identified MTB strains compared to other non-CaCO_3_ storing MTB strains, and the intricate interplay between MTB and other microorganisms in microbial communities within these ecosystems.

**Fig. 4.**
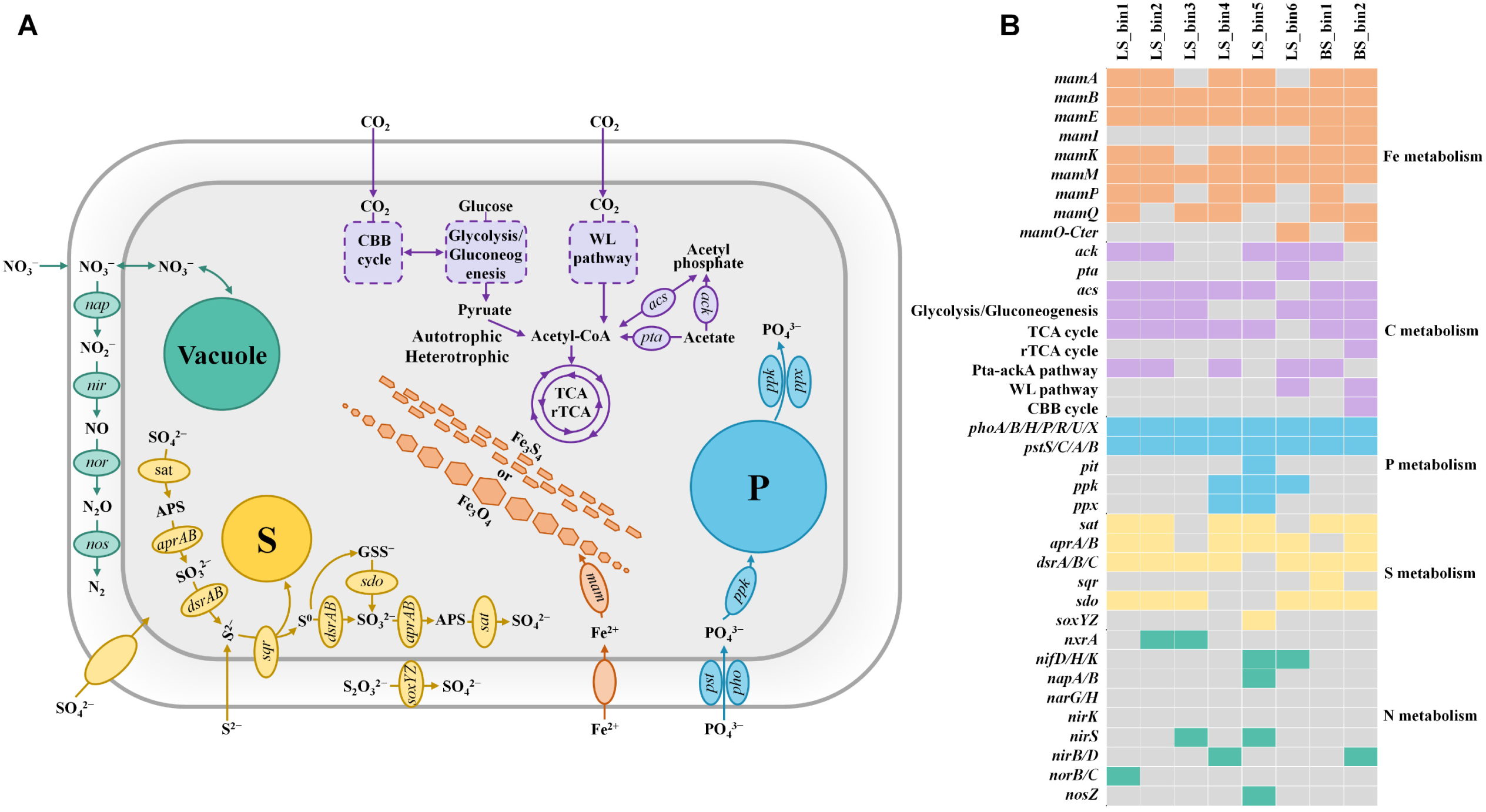
(A) Schematic diagram of metabolic pathways of MTB proposed based on metagenomic analysis and STEM images. Genomes were recovered from magnetically enriched MTB samples. The chosen metabolic pathways informed by the constructed genomes are: iron (Fe), carbon (C), phosphorus (P), sulfur (S), and nitrogen (N) metabolisms. (B) Genes involved in the biomineralization of the magnetosome, autotrophic and heterotrophic carbon utilization, sulfate reduction, sulfide oxidation, and nitrate reduction. Orange/purple/blue/yellow/green colors indicate the presence of genes, and gray represents the absence of genes. BS_bin1 and BS_bin2 were identified in both Flensborg Fjord and Faxe Bugt, while LS_bin1–bin4 and LS_bin5–bin6 were obtained in Östra Silen and Furesø, respectively (Supplementary Information Fig. S5).

**Fig. 5.**
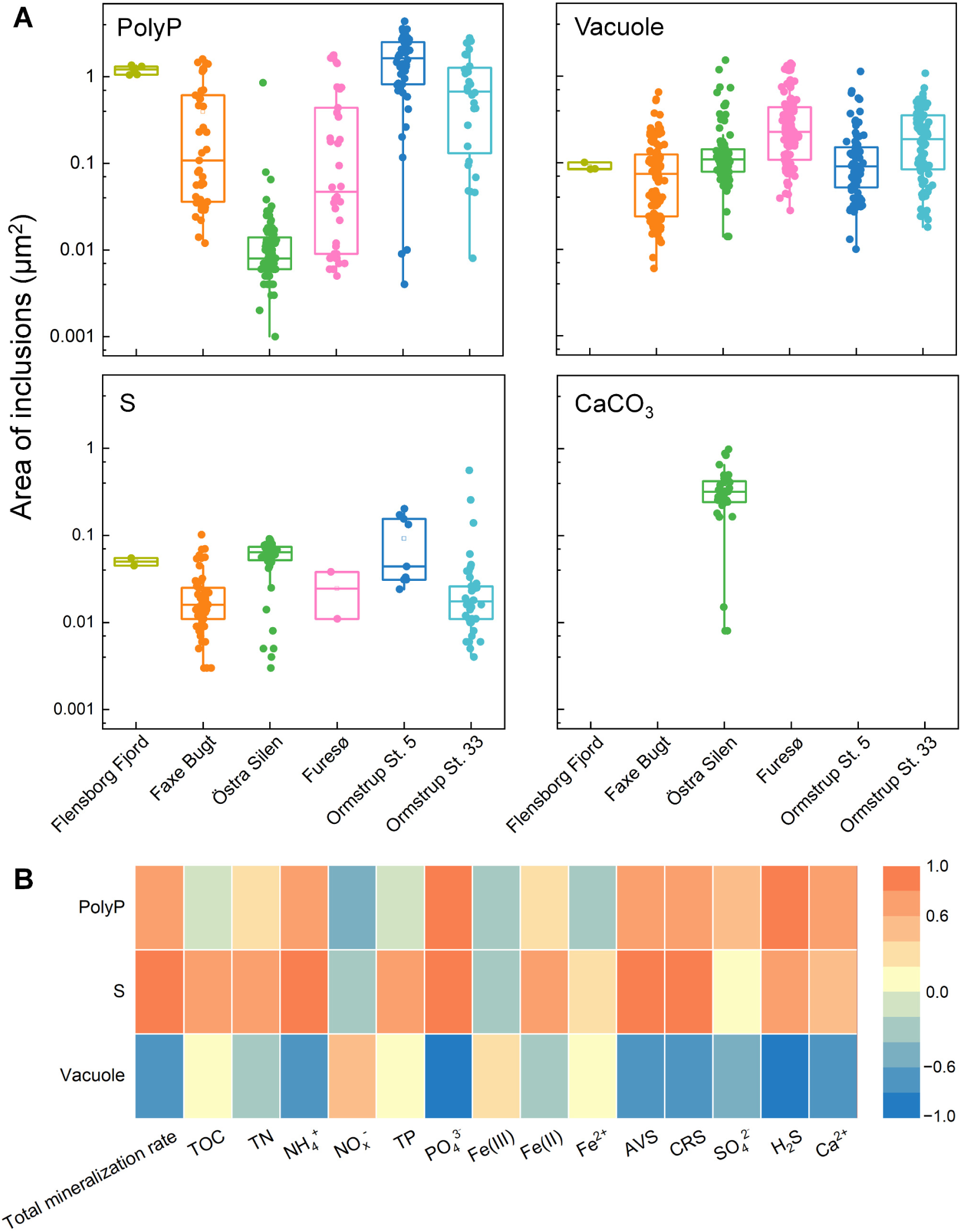
(A) Area of polyphosphate (polyP) inclusions, calcium carbonate (CaCO_3_) granules, sulfur (S) globules, and vacuoles in MTB cells (n = 48–470). (B) Correlation of the area of polyP inclusions, S globules, and vacuoles with depth-integrated values of geochemical parameters. A correlation coefficient greater than 0.6 or less than −0.6 indicates a strong positive or negative correlation, respectively.

### PolyP inclusions

The most abundant type of inclusions identified in MTB cells was the P-rich granules, which were observed in unicellular coccoid cells, rods, vibrios, and spirilla across all our sampling sites (Fig. 2–3). The granules also contained O and magnesium (Mg) or Ca as main elements (Fig. 3, Fig. S6). A dark red-purple color was observed when magnetically collected MTB cells were stained with toluidine blue (Fig. S7). Both observations indicated the presence of inclusion polyP as the storage form of P. PolyP is generally composed of linear polymers of orthophosphate linked through high-energy phosphoanhydride bonds ^18,84^. Each orthophosphate unit carries a monovalent negative charge at physiological pH, resulting in a large cation exchange capacity of polyP. The binding energy facilitates polyP to sequester Mg^2+^, Ca^2+^, K^+^ etc., resulting in the observed high electron density of the granules ^18,85^.

In this study, we found polyP inclusions ranging in size from 0.042–3.7 µm in width and 0.0010–4.4 µm^2^ in area, occupying 0.18–95% of the cytoplasmic space (n = 231) (Fig. 5A). For instance, magnetotactic cocci typically contained 1–2 spherical P-rich granules with the area of 0.0080–3.6 µm^2^, occupying up to 95% of the cytoplasmic space (n = 71). Similar P-rich granules have previously been identified in MTB using analytical and staining methods, which were also interpreted as polyP ^15,17–21^. For instance, 52–94% of the total MTB population in the water column of Lake Pavin were found to contain polyP inclusions, with the granules occupying a significant part of the cytoplasmic space in magnetotactic cocci ^17,78^. Similarly, Ca- and Mg-rich polyP inclusions have been found in MTB affiliated to the *Magnetococcus* genus in the suboxic zone of the Black Sea ^20^, in strain FCR-1 affiliated to the novel genus *Ca.* Magnetaquiglobus chichijimensis isolated from Chichijima, Japan ^86^, in *Ca.* Magnetoglobus multicellularis in Araruama Lagoon sediments ^21^, as well as in the magnetotactic spirillum strain XQGS-1 obtained from freshwater sediments in China ^27^.

The characterized polyP inclusions were supported by functional annotation of genes encoding for pathways associated with dissolved PO_4_^3−^ assimilation into polyP in some of the draft genomes (Fig. 4). PolyP-accumulating microorganisms generally take up PO_4_^3−^ by low-affinity PO_4_^3−^ transporters (e.g., Pit) and/or by high-affinity ABC-type PO_4_ ^3−^-specific transporter (Pst) ^22,87^. While genes encoding for Pit transporters were only identified in LS_bin5 (*Magnetospirillum*) retrieved from the sediment in Furesø, all our constructed genomes contained the genes coding for the Pst transporter (*pstS*, *C*, *A*, *B*) in addition to the PO_4_^3−^ uptake transcriptional regulator (*phoA*, *B*, *H*, *P*, *R*, *U*, *X*). The Pst transporter and Pho regulon have been described in the previous full-genome studies of the magnetotactic cocci strain MC-1 and the *Magnetospirillum* strains AMB-1, MS-1, and MSR-1 ^22,87,88^. PolyP kinase encoded genes *ppk* were identified in two *Magnetospirillum*-affiliated genomes, i.e., LS_bin4 from Östra Silen and LS_bin5 from Furesø, as well as in one *Desulfamplus*-affiliated genome LS_bin6 from Furesø. PolyP kinase is a principal enzyme that reversibly catalyzes the transfer of the terminal phosphate from ATP (PPK1) or GTP (PPK2) to an active site on the protein, initiating the processive synthesis of a polyP chain ^24,87^. PolyP-accumulating microorganisms have been suggested to evolve several mechanisms for efficient P utilization ^24,89^. Exopolyphophatase, encoded by *ppx*, is involved in degrading polyP into smaller branches of PO_4_^3–24^. Intriguingly, no *ppk* and *ppx* genes were found in the draft genomes affiliated with *Magnetococcia* (LS_bin1–3 and BS_bin1) and *Desulfobacteria* class (BS_bin2) (Fig. 4). This is in contrast to previous studies, where the presence of PPK and PPX was demonstrated in genomes of *Magnetococcales* (e.g., strain MC-1) and *Desulfobacterales* (e.g., BW-1 and HK-1) ^20,22^. We cannot rule out the fact that the corresponding genes were not covered by the sequencing. In continuation of this, it has been postulated that the ability to synthesize and degrade polyP in MTB might not be limited to PPK- and PPX-like enzymes ^22^. Genes encoding the enzyme polyphosphate: AMP phosphotransferase PAP, catalyzing the transfer of the terminal PO_4_^3−^from polyP to AMP and subsequently to ADP, also appeared to be absent in our eight draft genomes.

The physiological function of polyP in MTB remains unclear. PolyP may serve multiple functions, including motility, stress adaptation, buoyancy control, energy management, maintenance, and growth ^24^. In studies of different bacteria with the ability to accumulate polyP, such as PAO in EBPR reactors of wastewater treatment trains, it has been proposed that the storage of polyP inclusions serves as a source of ATP or a response to oxidative stress ^90^. The area of polyP inclusions per cell was positively correlated with depth-integrated porewater concentrations of PO_4_^3^⁻, Ca^2+^, NH_4_^+^, and H_2_S (Fig. 5B). Strong dependence of polyP synthesis on the PO_4_^3^⁻ availability has been defined in polyP-accumulating microorganisms such as PAO in engineered EBPR systems and sulfur-oxidizing bacteria in natural environments ^31,91,92^. Under excess PO_4_^3^⁻, luxury uptake occurs through the accumulation of polyP beyond its immediate metabolic needs for future use when PO_4_^3^⁻ becomes limited ^91,92^. Positive correlation of polyP inclusion size with porewater Ca^2+^ is consistent with the fact that Ca^2+^ is the major physiological divalent cation for polyP (Fig. 3, Fig. 5B, Fig. S6) ^18,85^. The positive correlation between the area of polyP inclusions per cell and H_2_S concentrations contrasts with the observation by Rivas-Lamelo et al and Schulz-Vogt et al ^17,20^. The authors hypothesized that magnetotactic cocci accumulate high amounts of PO_4_^3−^ in the form of polyP under oxic conditions, while polyP is degraded with the subsequent release of PO_4_^3−^ under sulfidic conditions ^17,20^. In addition, Rivas-Lamelo and coworkers (2017) speculated that the massive accumulation of polyP in magnetotactic cocci in Lake Pavin was due to the effect of oxic-anoxic fluctuations, either by traveling vertically over short distances between the anoxic to oxic zone, or by being affected by seasonal depth shifts of the oxic-anoxic boundary. This hypothesis on polyP serving as an energy source for maintenance and growth was confirmed by batch incubations with the *Magnetospirillum* strain AMB-1 under oxic, anoxic, and transient oxic-anoxic conditions ^93^. Su and coworkers found that the biosynthesis of polyP was promoted during the growth of AMB-1 under oxic conditions, while polyP granules disappeared and growth ceased under anoxic conditions ^93^.

### S globules

Analyses with STEM-energy dispersive X-ray spectroscopy (EDX) identified another type of intracellular inclusions, which was characterized by a smaller size and lower electron density than the polyP granules and composed exclusively of elemental sulfur (S^0^) (Fig. 2–3). The S-rich globules ranging from 0.048–0.94 µm in width and 0.0030–0.56 µm^2^ in area (n = 151) were observed at all sampling sites (Fig. 5A). Magnetotactic cocci commonly contained 1–2 spherical S-rich globules, occupying 0.24–14% of the cytoplasmic space (n = 71) (e.g., Fig. 2 B6, D5, E1). A single rod-shaped MTB in Östra Silen had 47 S-rich globules, making up 51% of the cytoplasmic space (Fig. 2 C5). The observed S globules were similar with those reported in numerous uncultivated MTB in freshwater and marine habitats ^10,18,94,95^. For instance, Li and coworkers investigated the S globules in *Ca.* Magnetobacterium casensis using STXM coupled with Near Edge X-ray Absorption Fine Structure (NEXAFS) analysis, and found that the S globules were comprised of a linear polymeric sulfur structure (S_5_^+^ and S_3_^+^) with the degree of polymerization increasing toward the core of S globule ^13^. Meanwhile, S globules in *Ca.* Magnetobacterium bavaricum were interpreted as cyclo-octasulphur (S_8_) using confocal Raman micro-spectrometry ^6,96^.

Genes related to sulfur oxidation and SO_4_^2−^ reduction were present in all the constructed MTB genomes, independent of the differences in SO_4_^2−^ and H_2_S concentrations found at the different sites (Fig. 4, Table S2). Looking into the genomic data of our assembled MAGs, the complete set of genes involved in dissimilatory SO_4_^2−^ reduction, including *sat*, *apr*, and *dsr* were found in LS_bin1, LS_bin2 (*Magnetococcia*), LS_bin4 (*Magnetospirillum*), and BS_bin2 (*Ca.* Magnetomorum), while the remaining constructed genomes contained only partial genes associated with this pathway (Fig. 4, Table S5). Moreover, the same enzymes (i.e., SAT, APR, and DSR) could operate in the reverse direction, mediating the oxidation of S^2−^, S^0^, and sulfite (SO_3_^2−^) to conserve energy for cell growth ^97^. For instance, many Nitrospirota MTB encode a complete or partial set of genes for both dissimilatory sulfur reduction and sulfur oxidation pathways ^13,98,99^. Consistent with our results for BS_bin2, the previous genomic prediction of *Ca.* Magnetobacterium casensis indicated that cells have the potential to reduce SO_4_^2−^ to S^2−^and oxidize S^2−^ to SO_4_^2−^, enabling them to switch between oxidation and reduction depending on environmental redox conditions ^13^. The presence of genes coding for [FeFe]- and [NiFe]-hydrogenase in BS_bin2 suggested that it may be capable of autotrophic growth using SO_4_^2−^reduction coupled with hydrogen metabolism, similar to SO_4_^2−^ reducing bacteria *Desulfovibri* ^100,101^. We also identified gene *sqr* encoding for sulfide:quinone oxidoreductase, which catalyzes the oxidation of S^2−^ to S^0^ and lead to the formation of glutathione persulfide (GSSH) as well as gene *sd*o encoding for sulfur dioxygenase, which oxidizes sulfane sulfur in GSSH to SO_3_^2−^, in several constructed genomes (Fig. 4). Along with APR and SAT, the oxidation of S^2−^may proceed all the way to SO_4_^2−^. The subunits of the periplasmic sulfur-oxidizing enzyme complex, coded by *soxYZ*, were found in the genomes of LS_bin5 (*Magnetospirillum*) and BS_bin1 (*Magnetococcia)*, which catalyze the oxidation of S_2_O_3_^2−^ to SO_4_^2−^. Genes for flavocytochrome c sulfide dehydrogenase (FccAB) that catalyze the oxidation of H_2_S to S^0^, transferring electrons to cytochrome c in the electron transport chain, were not identified in the constructed genomes. Collectively, the widespread distribution of S metabolic genes in MTB genomes indicates that they can adapt to local redox gradients by combining multiple S metabolic strategies. Their flexible metabolisms allow them to conserve energy through either sulfide oxidation or SO_4_^2−^ reduction as conditions dictate. They grow autotrophically by exploiting reduced sulfur compounds or hydrogen as energy sources.

Sulfur stored in globules is generally formed during the oxidation of reduced sulfur species: (i) sulfide oxidation catalyzed by SQR or FccAB, which produce polysulfides as intermediates that may contribute to the formation of intracellular sulfur globules; (ii) S_2_O_3_^2−^ oxidation catalyzed by the periplasmic SOX, which oxidizes S_2_O_3_^2−^ to store sulfane sulfur in S globules while the sulfone sulfur is hydrolytically released as SO_4_^2−^^102,103^. Our results, alongside previous studies, demonstrated the presence of sulfur globules and the genetic potential for oxidizing reduced sulfur compounds in diverse MTB from both freshwater and marine habitats ^13,16,22,25,87,98^. The identification of S globules in many uncultured magnetotactic cocci has been reported in environments with both high and very low concentrations of H_2_S, suggesting a chemolithoautotrophic or mixotrophic metabolism based on the oxidation of reduced sulfur compounds (Fig. 4) ^18,87^. S^0^ in globules serves as an energy storage that could be oxidized to SO_3_^2−^ by reverse-acting DsrAB, and further to SO_4_^2−^ by reversed SAT, as seen in the mixotrophic sulfur oxidizers like *Thioploca*, or reduced to sulfide as observed in the chemolithotrophic sulfur oxidizers like *Beggiatoa* ^33,104^. For MTB that lack enzymes to oxidize the stored S^0^, as in the case of LS_bin3, they may switch to anaerobic respiration under anoxic conditions, in which intracellular S^0^ serves as an electron acceptor perhaps with organic carbon acting as an electron donor, or via anaerobic S^0^ disproportionation, leading to production of S^2−^and SO_4_^2−^^105^.

### NO3^−^ vacuoles

Unlike the relatively high electron density observed in polyP inclusions and S globules, the white inclusions appeared electron-lucent and lacked detectable elements such as C, O, P, S, Ca, Mg, and Fe, even when compared to the surrounding cytoplasm (e.g., Fig. 2 F1–F6, Fig. 3). The observed white inclusions varied in number from 1 to 18 per cell and in size from 0.0060 to 1.5 µm^2^ (n = 470) (Fig. 5A). Morphological and chemical features of these white electron-lucent inclusions were identical to those widely discovered in MTB, such as *Ca.* Magnetobacterium casensis and the magnetotactic spirillum strain WYHS-1, where they were identified as NO_3_^−^-storing vacuoles ^13,15^. A large number of morphologically distinct MTB with vacuoles has been observed throughout the water column in Lake Pavin ^78^. The same type of white inclusions appeared in *Magnetospirillum magneticum* strain AMB-1, *Magnetospirillum gryphiswaldense* strain MSR-1, and *Ca.* Magnetoglobus multicellularis, where they were identified as PHA inclusions ^21,93,106^. In contrast to our observations, EDX analysis of PHA inclusions indicated a high C content compared to the surrounding cytoplasm ^93^. Together with the lack of genes coding for the key enzymes involved in PHA synthesis (e.g., β-ketothiolase (*phaA*), acetoacetyl CoA reductase (*phaB*), and PHA polymerase (*phaC*)) in the assembled genomes (data not shown) ^107^, the observed white inclusions were most likely vacuoles for NO_3_^−^ storage. Further research is essential to deepen our understanding of the nature and function of vacuoles in MTB. ^32,33^. Looking into the genomic data of our assembled MAGs, LS_bin5, affiliated with *Magnetospirillum* from Furesø, possessed the genomic potential for the denitrification pathway, as it contained most of the genes *napA/B*, *nirS*, and *nosZ* (Fig. 4, Table S5). In some *Magnetospirillum* species, periplasmic nitrate reductase encoded by *nap* has been shown to be involved in magnetite magnetosome biomineralization ^108^. The ability for incomplete denitrification was found in the draft genomes LS_bin3 and LS_bin1 (both *Magnetococcia*), where the reduction of NO_2_^−^ to nitric oxide (NO) by *nirS* and NO to nitrous oxide (N_2_O) by *norBC* was possible. Microbes with truncated incomplete denitrification pathways are well known ^13,20,76,109^. For example, Schübbe and colleagues described *napCBHGADF* and *norCBQ* in magnetotactic coccus strain MC-1 ^87^, while *narGHI* and *nosZ* were present in the constructed genomes affiliated to the phylum Nitrospirota ^110^.

We frequently observed that NO_3_^−^-storing vacuoles often coexisted with S globules in MTB cells (Fig. 2). Taken together with the putative NO_3_^−^-reducing and sulfur-oxidizing metabolisms identified in constructed MTB genomes (Fig. 4), these two metabolic processes were likely coupled in MTB cells. Coupled sulfide oxidation and denitrification have previously been suggested based on the observation of intracellular S granules and NO_3_^−^-storing vacuoles in *Ca.* Magnetobacterium casensis and within Nitrospirota ^13,15^. The area of S globules per cell and depth-integrated porewater concentrations of PO_4_^3^⁻ and NH_4_^+^ were strongly and positively correlated, while the area of S globules per cell and SO_4_^2−^ and H_2_S concentrations showed a moderate positive correlation (Fig. 5B). In contrast, the area of NO_3_^−^-containing vacuoles showed a strong negative correlation with PO_4_^3^⁻, Ca^2+^, NH_4_^+^, and H_2_S. The nature and function of S globules and NO_3_^−^ vacuoles in MTB are largely unknown, and the direct effect of environmental parameters, including NO_3_^−^ and H_2_S, on the storage of these intracellular inclusions has not yet been examined. Chemotaxis experiments of model S- and NO_3_^−^-storing benthic sulfur bacteria revealed a negative correlation of intracellular NO_3_^−^ levels in filamentous *Thioploca* with H_2_S concentrations ^33^. This observation was attributed to the adaptive strategy of sulfur bacteria, which optimizes their access to both NO_3_^−^ and H_2_S via chemotaxis-mediated migration: They fill the vacuoles with NO_3_^−^ at the sediment surface, and then migrate into deeper layers, where they oxidize H_2_S to S^0^ and SO_4_^2−^ until the depletion of intracellular NO_3_^−^, triggering an upward migration ^33,111^. Thus, similar to sulfur bacteria, the availability and distribution of sulfur compounds and NO_3_^−^ could drive the synthesis and degradation of inclusions in MTB.

### The ability of MTB to thrive in different environments

Of the 118 examined MTB cells, polyP granules (75%) and NO_3_^−^-containing vacuoles (67%) were the most common inclusions followed by S globules (25%) and CaCO_3_ granules (8%), while 15% of the MTB cells did not contain any inclusions (Fig. 5A). Of the polyP-containing cells, the occurrence of polyP frequently coincided with NO_3_^−^-containing vacuoles (72% of cells) or S globule (23% of cells), while 20% of examined cells contained all three inclusions. More than 10% of the examined cells contained both S globules and NO_3_^−^-containing vacuoles, but lacked polyP inclusions. The presence of MTB containing all polyP inclusions, S globules, and NO_3_^−^ vacuoles was observed at four sampling sites, except Faxe Bugt, despite the distinct physical-chemical and microbiological characteristics of the four sites. Together with the observed distribution of MTB in both oxic and anoxic zones, and the metabolic potential revealed by our assembled MTB genomes (Fig. 1, 4), our findings highlight their ability to switch between different redox-related metabolic activities, underscoring their high environmental adaptability and resilience.

To further illustrate this, we predicted the possible metabolisms of MTB cells connected to S-, polyP- and NO_3_^−^-rich inclusions, based on our results and relevant literature (Fig. 4, 6). The observed presence of S-rich inclusions together with widespread distribution of S metabolic genes indicate that both marine and freshwater MTB resemble sulfur bacteria, like BS_bin2 and LS_bin6 (Fig. 4, Fig. S5). In the oxic regions of the sediments, the stored elemental S in the S globules is acting as an electron donor, using O_2_, as the electron acceptor (Fig. 6). The associated energy is used to support cell growth and/or to sequestrate PO_4_ ^3−^ and/or NO_3_^−^ stored in intracellular polyP inclusions and NO_3_^−^ vacuoles ^9,10,93^. Using geomagnetic navigation, MTB may shuttle downwards into the nitrogenous zone to reduce bulk NO_3_^−^, by using organic carbon as an electron donor or the stored S^0^, as either mixotrophic or strict chemoautolithotrophic MTB (Fig. 6). The gained energy is utilized for growth and/or assimilation of NO_3_^−^ beyond their metabolic needs and polyP storage. When moving further downwards to the reduced zone, bulk H_2_S is taken up and oxidized to S^0^, which is stored. The oxidation of sulfide is coupled to the reduction of NO_3_^−^ stored in vacuoles during chemolithoautotrophic growth (Fig. 6) ^13,16,112^. Some MTB might also switch to heterotrophic metabolism through dissimilatory SO_4_^2−^reduction or disproportionation of elemental sulfur ^13,16,105^. After refilling the S globules in the reduced zone, MTB may shuttle back to the upper oxic/nitrogenous region along the geomagnetic field lines in their search for preferable electron donors.

**Fig. 6.**
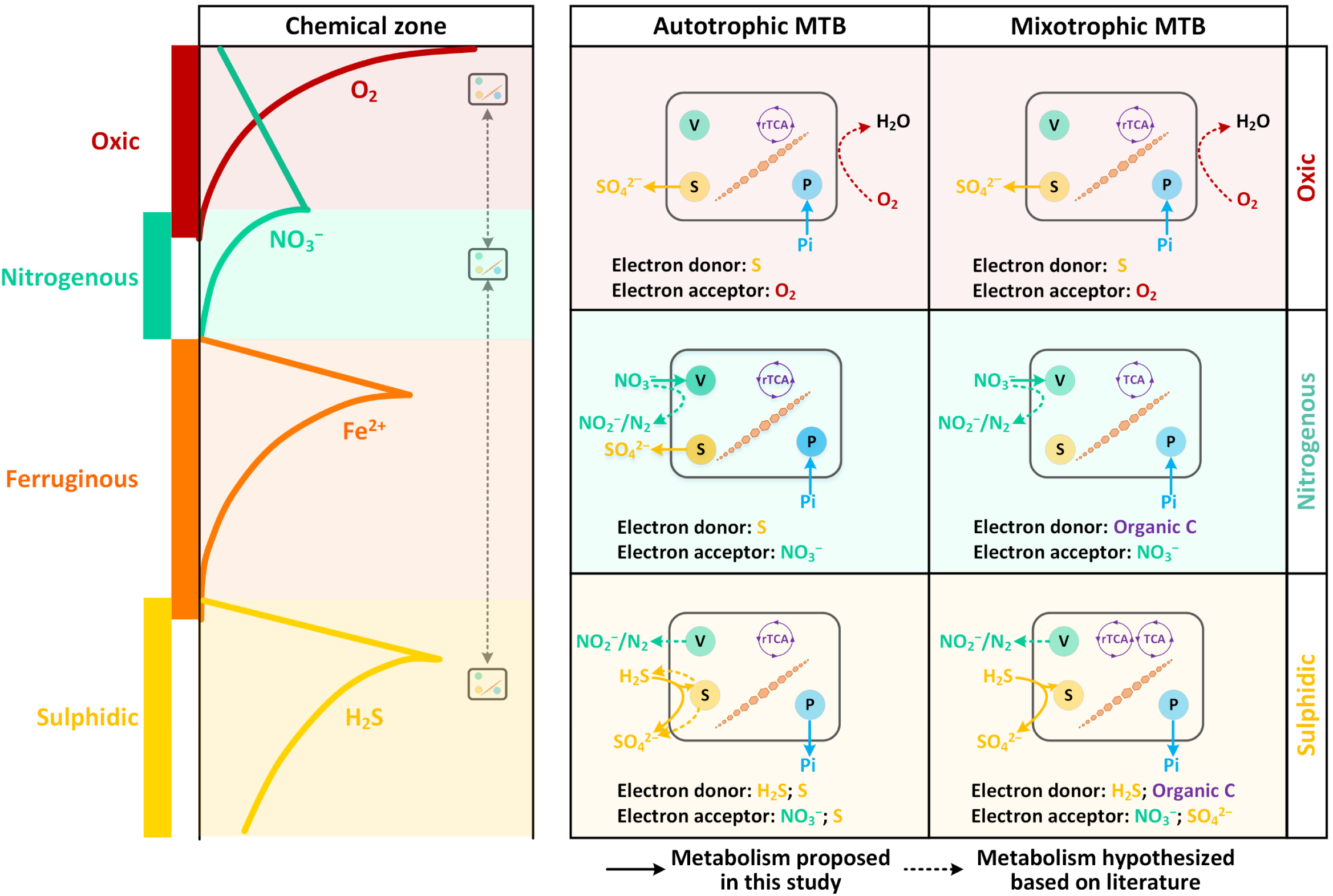
Proposed synthesis and utilization of sulfur globules, nitrate-storing vacuoles, and polyphosphate inclusions by MTB across idealized zonation in the sediment. The sulfidic zone does not always exist in freshwater sediments. The metabolisms related to intracellular inclusions of MTB are proposed based on microscopic, staining, and metagenomic analyses in this study and the literature.

### Environmental implications of P-, N-, and S-rich inclusions

In order to access the quantitative impact of MTB inclusions in the freshwater sediments of Östra Silen, Furesø, and Lake Ormstrup, the following assumptions are made: (i) The numbers of MTB cells range between 10^3^ and 10^6^ cells/mL in sediments ^10,55,95^; (ii) MTB are found in the upper 8 cm of sediments; (iii) The generation time for all freshwater MTB is approximately 12 h ^57^; (iv) Lakes are considered as “closed systems”. The latter is especially true for Lake Ormstrup, which is characterized by a very high hydraulic retention time of > 1 year ^40^. Given the average numbers and size of intracellular inclusions in MTB characterized in this study, our conservative calculations showed that approximately 1.8 × (10^−12^–10^−9^) kg P/cm^2^, 4.8 × (10^−12^−10^−9^) kg N/cm^2^, and 7.3 × (10^−13^−10^−10^) kg S/cm^2^ are stored within MTB cells (see Table S6–S7 for calculation details). When extrapolated to an annual scale, these numbers corresponded to 6.6 × (10^−10^–10^−7^) kg P/cm^2^, 1.8 × (10^−9^–10^−3^) kg N/cm^2^, and 2.7 × (10^−10^–10^−7^) kg S/cm^2^, respectively. To contextualize these findings to our study sites in Östra Silen, Furesø, and Lake Ormstrup, the intracellular P, N, and S stored in MTB cells accounted for at least 0.60%, 0.91%, and 0.23% of the total sedimentary P, N and S in the upper 8 cm, respectively (Table S6). Notably, the calculations do not capture the turnover of polyP, NO_3_^−^, and S inclusions in MTB. Nevertheless, our estimates underline the important role of MTB in the biogeochemical cycles. MTB associated transport of PO_4_^3−^ and NO_3_^−^ from the surface into deeper layers can have important implications for the preservation of P and N, reducing algae growth in the surface waters. In practice, high storage capacity of MTB for PO_4_^3−^ and NO_3_^−^offers a sustainable approach for nutrient recovery from water.

## Materials and Methods

### Study sites and sampling

In this study, sediments were sampled at five different sites, including two brackish sites (i.e., Flensborg Fjord and Faxe Bugt) and three freshwater lakes (i.e., Östra Silen, Furesø, and Lake Ormstrup) (Fig. S1, Table S1). Flensborg Fjord is a eutrophic threshold fjord between Germany and Denmark ^113^. Our sampling site is located at the shore near the sill, separating the inner and outer parts of the fjord. Faxe Bugt is a bay in the western part of the Baltic Sea. Also, the sampling site in Faxe Bugt is placed at the shore. Furesø is large mesotrophic lake (surface area: 9.4 km^2^) located ∼20 km north of Copenhagen on the island of Zealand ^114^. The mean hydraulic retention time is ∼10 years. The lake is affected by eutrophication through domestic waste from 1900 to 1970, where sedimentation of phosphorus and nitrogen increased in the lake. The phosphorus content reached up to 1 mg/g DW in the shallow surface sediment^115^. The sampling site in Östra Silen, Sweden, is situated on the shore in the north-eastern part of the lake, which receives the main inflow from the River Göta. As a drinking water source and fish reproduction site, Östra Silen is relatively pristine without regular nutrient input, as the lake catchment is dominated by coniferous forests. The sediments in Flensborg Fjord, Faxe Bugt, Östra Silen, and Furesø were well-sorted fine sand with a median grain size (Table S1). Lake Ormstrup is a small, shallow hypertrophic lake with an open surface area of 0.11 km^2^. The estimated hydraulic retention time is >1 year. Lake Ormstrup was characterized by soft muddy sediment with extremely high water content of 94–95% (Table S1). Sediment was collected at Station 5 and 33 in the western and eastern parts of the lake, respectively ^40,116^. Notably high TP concentrations, reaching up to 6 mg/g DW in the upper 5 cm of the sediment, have been detected in Lake Ormstrup, largely due to the historical release and feeding of ducks in the lake 30–40 years ago ^40^.

Sediment and water samples were collected from the five locations from July to September 2021 (Table S1). Temperature, salinity, and pH in the bottom water at each site were measured using probes (WTW GmbH, Weilheim, Germany). Sediment cores were retrieved by diving or using a Kajak corer (KC Denmark). Sediment cores were transported to the laboratory within 2–4 hours, stored in an aerated tank filled with water from the sampling sites, maintained at a temperature close to *in situ* conditions until further handling. The total carbon mineralization rate (mmol/m^2^/d) was determined the day after sampling by measuring the total benthic O₂ uptake rate of sediment cores (the analytical details were described in Supplementary Note).

### Solid-phase and porewater analyses

Sediment cores from each site were sliced at 1 cm intervals down to 8 cm depth under N_2_ atmosphere in a glove bag. The sediments from each depth interval were homogenized and subsampled for solid-phase analyses of grain-size, poorly crystalline Fe oxides and reduced Fe, AVS (iron sulfide (FeS) + H_2_S) and CRS (S^0^ + iron disulphide (FeS_2_)), TP, TOC, and TN. The details of sediment extraction and conservation and the measurement of these solid-phase parameters were listed in Supplementary Note. Additionally, ∼1 g of homogenized sediments was stored at –80 °C until DNA extractions. Subsamples of homogenized sediment from each depth interval were centrifuged (4000 rpm for 10 min) at 4 °C, and the supernatant was filtered through 0.22 µm-pore-diameter polypropylene filters under N_2_ atmosphere. The preservation and measurement of porewater samples for the different analyses were described in Supplementary Note.

### Magnetic enrichment of MTB

Magnetic enrichment of MTB was performed immediately upon return to the laboratory. The collected sediment from each site was homogenized and transferred to 500–1000 mL glass beakers with a ratio of sediment to on-site water of 2:1 ^56^. The concentrated cells in the MTB pellet were examined using the hanging drop technique under a light microscope ^117^. Evidence of magnetotaxis was confirmed by a rotating magnet. Sufficient living MTB cells were separated into two fractions for STEM-EDX analysis and DNA extraction.

### Scanning transmission electron microscopy and energy dispersive X-ray spectroscopy analysis

20 µL of magnetically collected MTB sample was loaded onto a 200-mesh Formvar-carbon-coated copper grid (Agar Scientific, USA). The cells were allowed to settle for 30 min, and excess water was carefully removed with filter paper (Whatman, Germany). The grid was left to dry before microscopic analysis. Cell visualization and elemental analysis were conducted in STEM mode at Fei Quanta FEG 200 ESEM, equipped with an Oxford Instruments EDX spectrometer. The electron microscope was operated at an accelerating voltage of 30 kV and a working distance of 10 mm. The diameter and area of MTB cells and inclusions were measured using ImageJ software (1.53v).

### 16S rRNA amplicon and shotgun sequencing

Extraction of DNA was performed on magnetically collected MTB samples and sediment from each individual 1-cm depth interval by using FastDNA™ SPIN Kit for Soil, following the manufacturer’s instructions (MP Biomedicals, USA). The quantity and quality of the extracted DNA were measured and checked by its 260/280 ratio with a NanoDrop (ThermoFisher Scientific, USA). For phylogenetic analysis, the extracted DNA was amplified using universal primers PRK341F (5’-CCTAYGGGRBGCASCAG-3’) and PRK806R (5’-GGACTACNNGGGTATCTAAT-3’) ^118^. Library preparation and Illumina HiSeq sequencing were carried out by DMAC (DTU Multi-Assay Core Facility, Denmark). For metagenomic analysis, shotgun DNA sequencing of magnetically collected MTB samples was performed as a 150 bp pair-end run on DNBSEQ-G400 at Beijing Genomics Institute’s (BGI) (Copenhagen, Denmark).

#### Phylogenetic analysis

Illumina data was analyzed in USEARCH to examine quality to optimize trimming procedures ^119^. Quality control, trimming, merging of paired ends, and error correction were performed in DADA2 ^120^. 16S rRNA genes were recovered from the reference genomes using Barrnap v0.9 (https://github.com/tseemann/barrnap). 16S rRNA gene sequences were aligned with MUSCLE in the MEGA X software using default settings ^121^. The maximum-likelihood tree was constructed using the Tamura-Nei model in MEGA X, and the tree was visualized with the online web tool from the Interactive Tree of Life (iTol) ^122^.

#### Metagenomic analysis

After quality trimming and filtering using SOAPnuke (v1.5.5) ^123^, the clean reads were assembled using SPAdes v3.15.3 ^124^. The assembled contigs were then grouped into bins using Vamb v3.0.6 Snakemake workflow ^125^ and binny v2.1.beta50-36 ^126^. The resulting bins were dereplicated with dRep v3.4.0 with average nucleotide identity (ANI) of 99% ^127^. Besides, the clean reads were co-assembled using megahit v1.2.9 ^128^, and subsequently sorted into bins using six different binning tools, including binny v2.1.beta50-36^126^, MetaWRAP v1.3.2 collection (MetaBAT2 and MaxBin2) ^129^, MetaBinner v1.4. ^130^, SemiBin v0.7.0 ^131^, and MetaDecoder v1.0.13 ^132^. The bins were produced from the top three binners that yielded the highest number of high-quality metagenomic assembled genomes (MAGs), and merged using the bin refinement module of MetaWRAP v1.3.2. Bins, generated from individual assembly and co-assembly, were combined for collective dereplication using dRep v3.4.0 with 99% and 95% ANI ^127^.

Completeness and contamination of bins were assessed using dRep v3.4.0, and only genomes with >70% completeness and < 5% contamination were retained. The identification, annotation, and visualization of magnetosome gene clusters (MGCs) were performed using MAGcluster with manual inspection ^133^. A total of eight MGC-containing genomes were recovered, among which five were high-quality and three were medium-quality (Table S4). Phylogenetic analyses of genomes were conducted with the GTDB-Tk v2.1.0 ^134^ using the de novo workflow with a set of 120 translated universal single-copy genes and the genome taxonomy database (GTDB) ^135^. The relative abundances of MTB MAGs in magnetically collected samples were quantified with CoverM v0.6.1 (https://github.com/wwood/CoverM). A curated set of genes involved in C, N, and S metabolism was searched using METABOLIC v4.0 ^136^, and genes involved in P metabolism were annotated through the well-curated P cycling database (PCycDB) v1.1 ^137^. To identify *mam* genes within reads, MAG sequences were first indexed using the bwa index, and clean reads were mapped to these indexed MAG sequences using bwa mem. The resulting unique BAM files were sorted and indexed and thereafter converted to FASTA format. The FASTA files were blasted with the *mam* gene databases using Diamond blastx, which allowed us to identify all potential *mam* genes. The quantification of *mam* genes with sediment depths was determined by dividing the number of *mam* genes by the number of clean reads in each sediment sample.

### Statistical correlation analysis

All statistical tests were performed using R v3.3.1. Pearson correlation analysis was performed to evaluate the correlations between different geochemical parameters. The Mantel test was conducted to examine the correlation of the depth distribution of the *mam* gene in sediments with geochemical parameters.

### Data availability

16S rRNA gene sequences amplified from magnetically collected MTB samples and homogenized sediments from each depth interval are available in NCBI under accession number PRJNA1013417. The constructed MTB genomes were deposited in NCBI under the accession number PRJNA1013564.

## Acknowledgments

The work was supported financially by the National Natural Science Foundation of China (grant NO. 52300028) and the VILLUM FONDEN (grant NO. 00023110). M.M.J. and I.A. were supported by the ERC-advanced grant 101098064–ANAEROB. This work has been performed using the Danish National Life Science Supercomputing Center, Computerome. The authors thank Heidi Grøn Jensen and Anna-Marie Klamt from the University of Southern Denmark for their assistance during solid-phase and porewater analyses. We also thank Iben Kjær Bock for her help during sampling in Östra Silen.

