## Supplementary Information for "Beyond magnetosomes: ubiquitous and diverse intracellular inclusions expand the role of magnetotactic bacteria in biogeochemical cycling"

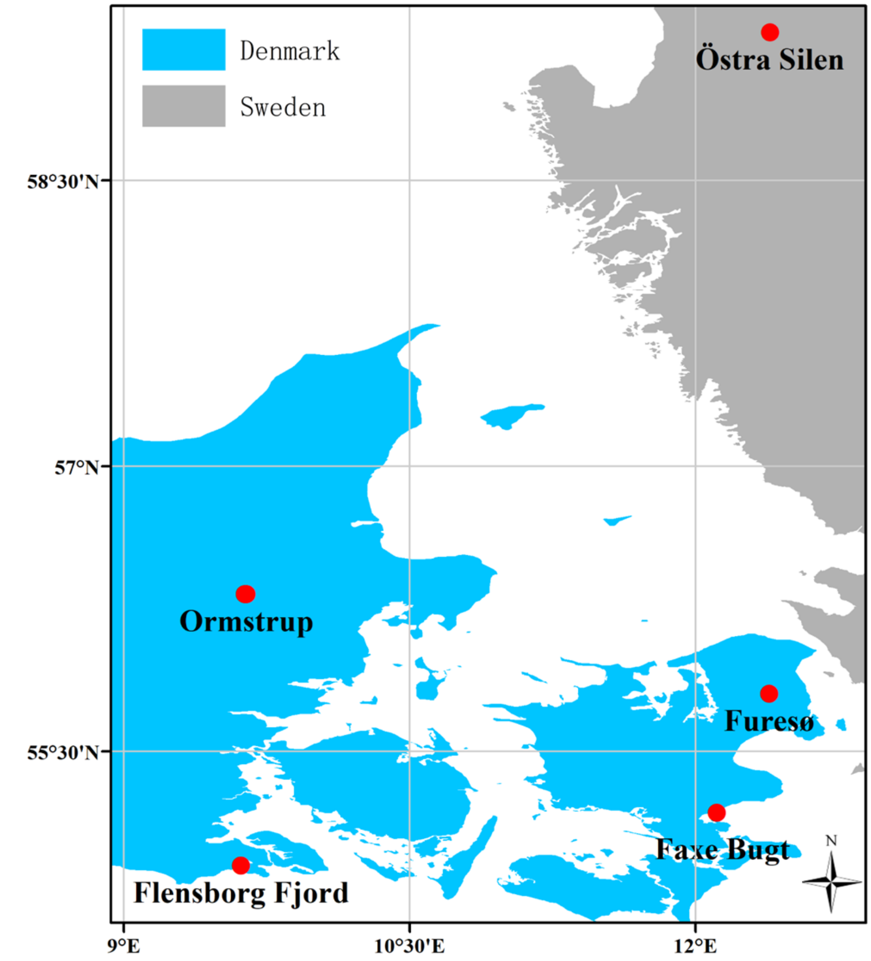


Fig. S1. Map of the sampling sites.

Table S1. General characteristics of sampling sites. Values of median grain size, silt-clay content, porosity, density, and water content were averaged over the 0 to 8 cm depth stratum.

| Sampling site | Lat., N | Long., E | Water depth (m) | Salinity (‰) | Temperature (˚C) | $O_2$ in the bottom water (µmol/L) | Median grain size (mm) | Silt-clay content (%) | Sediment type | Density ($g/cm^3$) | Porosity (v/v) | Water content (%) |
|---|---|---|---|---|---|---|---|---|---|---|---|---|
| Flensborg Fjord | 54°54'04" | 9°36'53" | 1.4 | 15 | 21 | 255 | 0.25 ± 0.005 | 0 | Fine sand | 1.9 ± 0.019 | 0.31 ± 0.012 | 17 ± 0.57 |
| Faxe Bugt | 55°10'42" | 12°06'40" | ~1.3 | 8 | 23 | 256 | 0.19 ± 0.026 | 0.13 ± 0.04 | Fine sand | 1.8 ± 0.050 | 0.30 ± 0.007 | 17 ± 0.65 |
| Östra Silen | 59°16'37" | 12°23'29" | 1.4 | 0 | 23 | 268 | 0.19 ± 0.012 | 1.9 ± 0.93 | Fine sand | 1.9 ± 0.064 | 0.36 ± 0.026 | 19 ± 0.82 |
| Furesø | 55°48'13" | 12°23'12" | ~1.2 | 0 | 18 | 296 | 0.29 ± 0.003 | 0 | Fine sand | 1.8 ± 0.037 | 0.30 ± 0.013 | 16 ± 0.47 |
| Ormstrup St. 5 | 56°19'33" | 9°38'10" | 1.8 | 0 | 14 | n.d. [a] | 0.26 ± 0.019 | 5.2 ± 0.61 | Muddy sand | 1.0 ± 0.009 | 1.0 ± 0.007 | 94 ± 0.60 |
| Ormstrup St. 33 | 56°19'35" | 9°38'37" | 4.5 | 0 | 14 | 4.9 | 0.22 ± 0.009 | 9.6 ± 0.41 | Muddy sand | 1.0 ± 0.004 | 1.0 ± 0.007 | 95 ± 0.44 |

[a] n.d. refers to the value not detected.

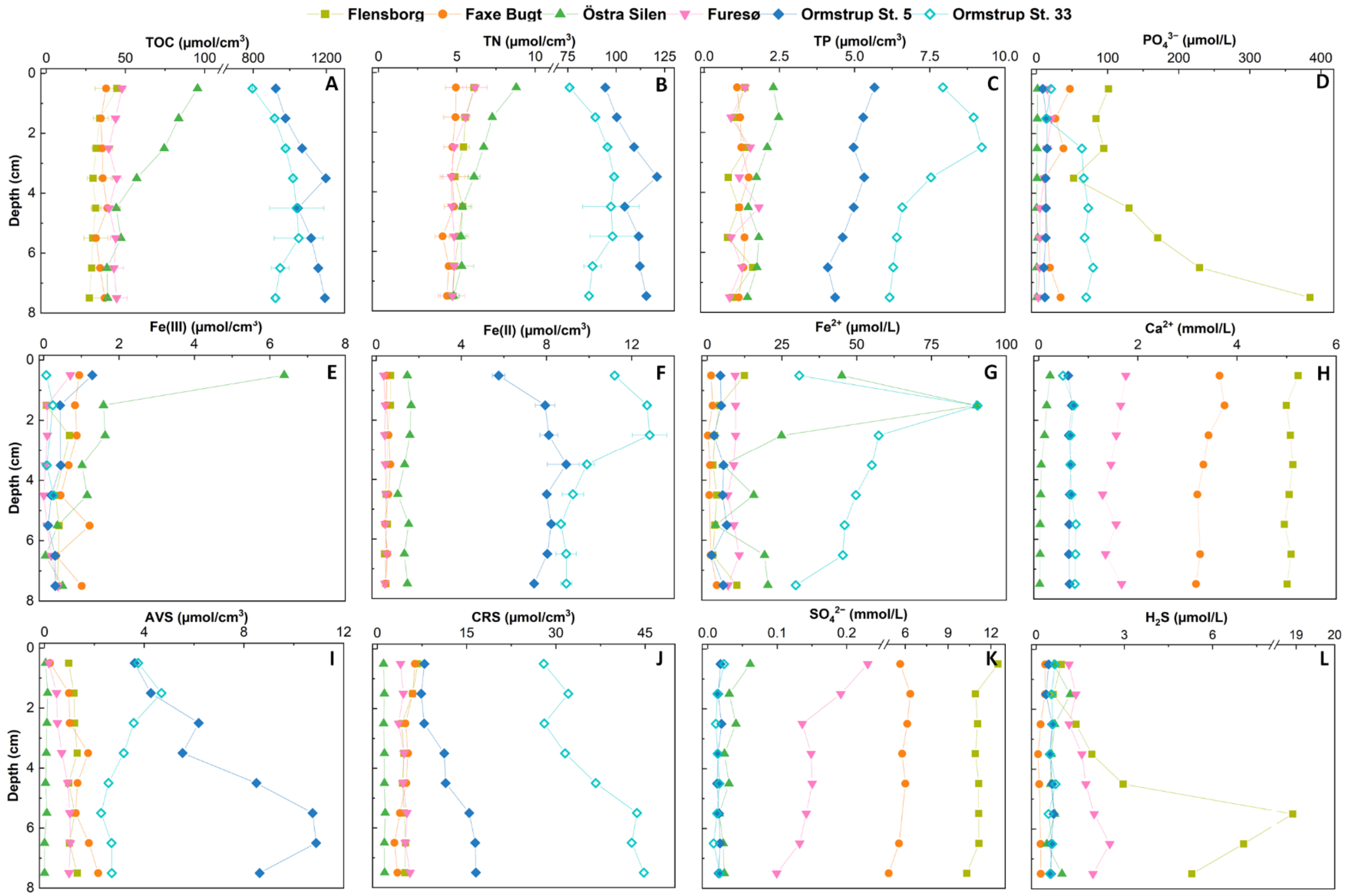


Fig. S2. Depth distributions of (A) total organic carbon (TOC), (B) total nitrogen (TN), (C) total phosphorus (TP), (D) $PO_4^{3-}$, (E) Fe(III), (F) Fe(II), (G) $Fe^{2+}$, (H) $Ca^{2+}$, (I) acid volatile sulfides (AVS), (J) chromium reducible sulfides (CRS), (K) $SO_4^{2-}$, and (L) $H_2S$ in Flensborg Fjord, Faxe Bugt, Östra Silen, Furesø, and Lake Ormstrup (Station 5 and 33). Error bars represent the range of duplicate measurements.

Table S2. Integrated values (mmol/$m^2$) of geochemical parameters. Values were integrated over 0 to 8 cm depth stratum.

| Sampling site | Total mineralization rate [a] | TOC [b] | TN [b] | $NH_4^+$ | $NO_x^-$ | TP | $PO_4^{3-}$ | Fe(III) | Fe(II) | $Fe^{2+}$ | AVS | CRS | $SO_4^{2-}$ | $H_2S$ | $Ca^{2+}$ |
|---|---|---|---|---|---|---|---|---|---|---|---|---|---|---|---|
| Flensborg Fjord | 32 ± 4.3 | 2559 ± 162 | 421 ± 38 | 12 | 0.0034 | 91 | 3.9 | 18 | 42 | 0.12 | 92 | 394 | 276 | 0.12 | 125 |
| Faxe Bugt | 12 ± 1.8 | 2841 ± 550 | 371 ± 46 | 6.6 | 0.14 | 100 | 0.56 | 63 | 42 | 0.033 | 105 | 371 | 138 | 0.040 | 71 |
| Östra Silen | 11 ± 1.1 | 4792 ± 89 | 496 ± 24 | 4.6 | 0.11 | 151 | 0.025 | 132 | 113 | 0.66 | 5.0 | 94 | 0.92 | 0.0041 | 3 |
| Furesø | 24 ± 9.3 | 3263 ± 442 | 379 ± 12 | 3.9 | 0.0088 | 98 | 0.24 | 16 | 34 | 0.21 | 59 | 358 | 3.7 | 0.019 | 37 |
| Ormstrup St. 5 | 40 ± 5.2 | 86740 ± 68 | 8686 ± 14 | 8.1 | 0.085 | 393 | 1.0 | 31 | 624 | 0.34 | 584 | 943 | 1.2 | 0.037 | 48 |
| Ormstrup St. 33 | 36 ± 13 | 76641 ± 3246 | 7289 ± 276 | 31 | n.d. [c] | 591 | 4.3 | 7 | 825 | 3.9 | 255 | 2875 | 1.1 | 0.041 | 51 |

[a] The unit of total mineralization rate is mmol/$m^2$/d. Error bars represent the standard errors of the slope of regression lines. [b] Error bars represent the range of duplicates. [c] n.d. refers to the value not detected.

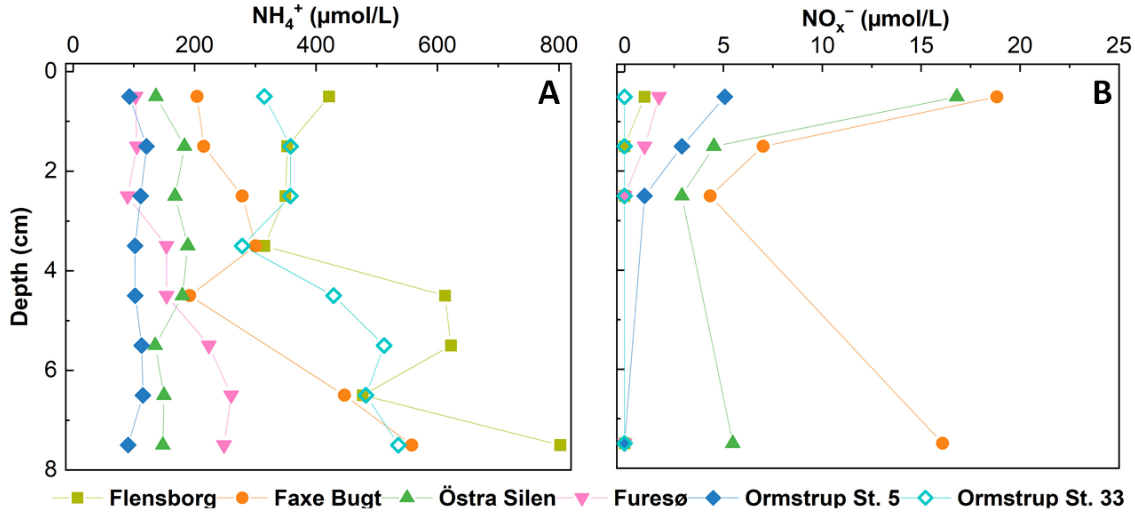


Fig. S3. Depth distributions of $NH_4^+$ and $NO_x^-$ ($NO_2^-$ + $NO_3^-$) concentrations in porewater.

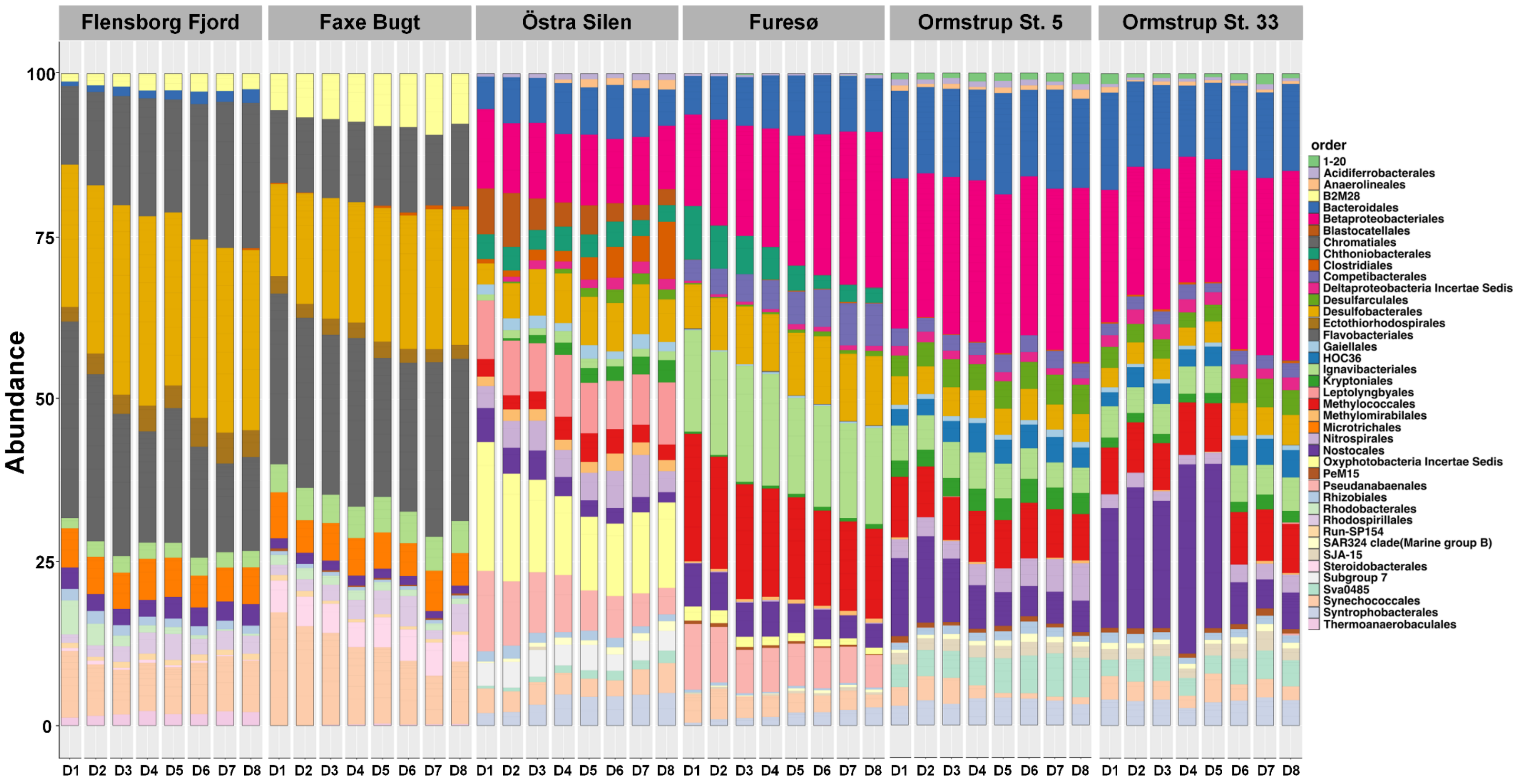


Fig. S4. Microbial community composition (at the order level) across sediment depths. Data was obtained from the 16S amplicon sequencing of the sediments in Flensborg Fjord, Faxe Bugt, Östra Silen, Furesø, and Lake Ormstrup (Station 5 and 33). Only the OTUs with relative abundance higher than 0.1% were represented.

Table S3. Size measurements of MTB cells and their intracellular inclusions. The diameter and area of MTB cells and inclusions were measured based on the scanning transmission electron microscopy images of MTB using ImageJ software.

| MTB ID [a] | Cell | | | | Intracellular inclusions | | | | | | | | | | | | | | | |
|---|---|---|---|---|---|---|---|---|---|---|---|---|---|---|---|---|---|---|---|---|
| | | | | | polyP granule | | | | S globule | | | | $CaCO_3$ granule | | | | $NO_3^-$-containing vacuoles | | | |
| | Shape | Length (µm) | Width (µm) | Area (µm$^2$) | Average diameter (µm) | Number | Average area (µm$^2$) | Area (%) | Average diameter (µm) | Number | Average area (µm$^2$) | Area (%) | Average diameter (µm) | Number | Average area (µm$^2$) | Area (%) | Average diameter (µm) | Number | Average area (µm$^2$) | Area (%) |
| A1 | Multicellular | 9.2 | 10 | 77 | / | / | / | / | / | / | / | / | / | / | / | / | / | / | / | / |
| A2 | Multicellular | 8.5 | 8.2 | 54 | / | / | / | / | / | / | / | / | / | / | / | / | / | / | / | / |
| A3 | Multicellular | 9.4 | 10 | 42 | / | / | / | / | / | / | / | / | / | / | / | / | / | / | / | / |
| A4 | Cocci | 2.3 | 1.8 | 3.4 | 1.4 ± 0.0030 | 2 | 1.26 ± 0.092 | 74 | / | / | / | / | / | / | / | / | 0.3 | 1 | 0.085 | 2.5 |
| B1 | Cocci | 3.1 | 2.7 | 7.4 | 0.22 ± 0.0070 | 2 | 0.031 ± 0.0015 | 0.82 | 0.36 ± 0.024 | 2 | 0.079 ± 0.023 | 2.1 | / | / | / | / | 0.43 ± 0.25 | 7 | 0.15 ± 0.12 | 14 |
| B2 | Cocci | 1.9 | 1.8 | 2.8 | 0.20 ± 0.029 | 2 | 0.029 ± 0.0065 | 2.1 | / | / | / | / | / | / | / | / | 0.33 ± 0.16 | 7 | 0.093 ± 0.073 | 24 |
| B3 | Cocci | 2.2 | 1.9 | 3.4 | 0.25 ± 0.022 | 2 | 0.043 ± 0.014 | 2.5 | 0.23 ± 0.061 | 3 | 0.038 ± 0.017 | 3.4 | / | / | / | / | / | / | / | / |
| B4 | Cocci | 2.2 | 21.0 | 3.4 | / | / | / | / | 0.17 ± 0.0020 | 2 | 0.024 ± 0.002 | 1.4 | / | / | / | / | 0.02 | 1 | 0.22 | 6.6 |
| B5 | Cocci | 1.8 | 1.4 | 2.0 | 0.73 ± 0.14 | 2 | 0.34 ± 0.11 | 34 | / | / | / | / | / | / | / | / | / | / | / | / |
| B6 | Cocci | 1.7 | 1.3 | 1.9 | 1.0 ± 0.015 | 2 | 0.70 ± 0.0080 | 74 | / | / | / | / | / | / | / | / | / | / | / | / |

| | | | | | | | | | | | | | | | | | | | | |
|---|---|---|---|---|---|---|---|---|---|---|---|---|---|---|---|---|---|---|---|---|
| B7 | Thumb-shaped | 2.0 | 1.4 | 2.5 | 0.69 ± 0.16 | 3 | 0.37 ± 0.17 | 45 | 0.21 | 1 | 0.03 | 1.2 | / | / | / | / | 0.37 ± 0.055 | 4 | 0.10 ± 0.025 | 16 |
| B8 | Rod | 2.9 | 1.0 | 1.9 | 0.94 ± 0.090 | 2 | 0.51 ± 0.046 | 54 | 0.16 | 1 | 0.015 | 0.8 | / | / | / | / | 0.20 ± 0.048 | 3 | 0.03 ± 0.010 | 4.6 |
| B9 | Vibrio | 3.3 | 0.9 | 2.1 | / | / | / | / | / | / | / | / | / | / | / | / | / | / | / | / |
| C1 | Rod | 11 | 4.4 | 36 | / | / | / | / | / | / | / | / | 3.3 ± 0.24 | 5 | 7.9 ± 1.8 | 100 | / | / | / | / |
| C2 | Rod | 1.7 | 1.0 | 1.3 | 0.2 | 1 | 0.029 | 2.2 | / | / | / | / | 0.86 ± 0.35 | 2 | 0.61 ± 0.37 | 94 | / | / | / | / |
| C3 | Rod | 9.6 | 6.0 | 52 | 1.2 | 1 | 0.85 | 1.6 | / | / | / | / | 4.8 ± 0.64 | 4 | 12 ± 1.9 | 90 | / | / | / | / |
| C4 | Rod | 2.9 | 1.8 | 4.8 | 0.30 ± 0.065 | 3 | 0.056 ± 0.023 | 3.6 | / | / | / | / | 0.64 ± 0.06 | 6 | 0.30 ± 0.16 | 38 | 1.0 ± 0.056 | 4 | 0.76 ± 0.27 | 64 |
| C5 | Rod | 4.5 | 1.6 | 5.4 | / | / | / | / | 0.26 ± 0.082 | 47 | 0.058 ± 0.023 | 51 | / | / | / | / | / | / | / | / |
| C6 | Rod | 5.5 | 3.5 | 16 | / | / | / | / | / | / | / | / | 0.69 ± 0.10 | 17 | 0.35 ± 0.09 | 38 | 1.3 ± 0.20 | 2 | 1.1 ± 0.40 | 14 |
| C7 | Rod | 1.7 | 1.1 | 1.6 | 0.077 ± 0.016 | 25 | 0.0050 ± 0.0015 | 7.8 | / | / | / | / | 0.12 ± 0.024 | 3 | 0.010 ± 0.0033 | 1.9 | 0.31 ± 0.080 | 12 | 0.07 ± 0.029 | 49 |
| C8 | Thumb-shaped | 2.8 | 3.6 | 1.7 | 0.095 ± 0.016 | 9 | 0.0073 ± 0.0019 | 3.9 | / | / | / | / | 0.62 ± 0.13 | 3 | 0.29 ± 0.08 | 24 | 0.33 ± 0.070 | 10 | 0.11 ± 0.031 | 63 |
| C9 | Rod | 5.6 | 3.6 | 1.8 | 0.13 ± 0.024 | 5 | 0.015 0.0041 ± | 4.1 | / | / | / | / | 1.0 ± 0.017 | 2 | 0.86 ± 0.02 | 94 | 0.34 ± 0.051 | 12 | 0.12 ± 0.024 | 79 |
| C10 | Rod | 10 | 2.4 | 19 | / | / | / | / | / | / | / | / | 1.7 ± 0.49 | 6 | 2.9 ± 1.2 | 95 | / | / | / | / |
| C11 | Rod | 6.7 | 2.1 | 11 | 0.11 ± 0.023 | 17 | 0.011 ± 0.0031 | 1.7 | / | / | / | / | / | / | / | / | 0.62 ± 0.057 | 3 | 0.31 ± 0.045 | 8.1 |
| C12 | Vibrio | 2.6 | 0.9 | 1.9 | / | / | / | / | / | / | / | / | / | / | / | / | 0.37 ± 0.060 | 8 | 0.10 ± 0.040 | 42 |

| | | | | | | | | | | | | | | | | | | | | |
|---|---|---|---|---|---|---|---|---|---|---|---|---|---|---|---|---|---|---|---|---|
| C13 | Thumb-shaped | 3.1 | 1.7 | 4.4 | 0.17 ± 0.014 | 5 | 0.022 ± 0.0042 | 2.5 | / | / | / | / | / | / | / | / | 0.37 ± 0.042 | 16 | 0.10 ± 0.030 | 37 |
| C14 | Rod | 4.1 | 1.3 | 4.2 | 0.20 ± 0.012 | 2 | 0.035 ± 0.0030 | 1.7 | / | / | / | / | / | / | / | / | 0.47 ± 0.16 | 17 | 0.18 ± 0.13 | 72 |
| D1 | Rod | 5.5 | 2.5 | 12 | 0.19 | 1 | 0.022 | 0.18 | / | / | / | / | / | / | / | / | / | / | / | / |
| D2 | Rod | 3.4 | 2.0 | 5.9 | / | / | / | / | / | / | / | / | / | / | / | / | 0.85 ± 0.16 | 8 | 0.40 ± 0.16 | 54 |
| D3 | Rod | 6.7 | 2.2 | 15 | / | / | / | / | / | / | / | / | / | / | / | / | 1.1 ± 0.32 | 11 | 0.91 ± 0.41 | 66 |
| D4 | Spirilla | 2.3 | 0.3 | 0.9 | 0.10 ± 0.0055 | 2 | 0.008 | 1.7 | / | / | / | / | / | / | / | / | / | / | / | / |
| D5-1 | Cocci | 2.6 | 1.9 | 4.6 | 1.4 ± 0.081 | 2 | 1.60 ± 0.15 | 69 | 0.12 | 1 | 0.011 | 0.24 | / | / | / | / | 0.32 ± 0.073 | 5 | 0.072 ± 0.027 | 7.8 |
| D5-2 | Cocci | 2.2 | 1.7 | 3.1 | 0.53 ± 0.0065 | 2 | 0.19 ± 0.0090 | 12 | 0.24 | 1 | 0.038 | 1.2 | / | / | / | / | / | / | / | / |
| D5-3 | Cocci | 2.4 | 2.3 | 4.4 | 0.20 ± 0.0035 | 2 | 0.040 ± 0.002 | 1.7 | / | / | / | / | / | / | / | / | 0.42 ± 0.12 | 13 | 0.17 ± 0.08 | 51 |
| E1 | Cocci | 1.8 | 1.4 | 2.1 | 0.92 ± 0.041 | 2 | 0.71 ± 0.055 | 67 | 0.2 | 1 | 0.031 | 1.5 | / | / | / | / | 0.31 ± 0.059 | 4 | 0.076 ± 0.025 | 14 |
| E2 | Cocci | 2.8 | 2.5 | 5.5 | 2.0 ± 0.064 | 2 | 2.5 ± 0.53 | 92 | 0.24 | 1 | 0.031 | 0.56 | / | / | / | / | 0.43 ± 0.027 | 3 | 0.13 ± 0.025 | 7.1 |
| E3 | Cocci | 3.6 | 2.4 | 7.7 | 2.2 ± 0.15 | 2 | 3.2 ± 0.34 | 83 | / | / | / | / | / | / | / | / | 0.45 ± 0.039 | 3 | 0.15 ± 0.009 | 5.7 |
| E4 | Cocci | 2.5 | 2.1 | 4.3 | 1.8 ± 0.096 | 2 | 1.8 ± 0.096 | 83 | 0.22 ± 0.010 | 2 | 0.034 ± 0.010 | 1.6 | / | / | / | / | 0.27 ± 0.021 | 2 | 0.08 ± 0.011 | 3.6 |
| E5 | Cocci | 2.7 | 2.6 | 5.6 | 1.7 ± 0.089 | 2 | 1.9 ± 0.14 | 69 | / | / | / | / | / | / | / | / | 0.43 ± 0.038 | 3 | 0.12 ± 0.019 | 6.6 |
| E6 | Cocci | 2.4 | 2.3 | 4.8 | 0.89 | 1 | 0.59 | 12 | 0.44 | 1 | 0.13 | 2.8 | / | / | / | / | 0.47 ± 0.31 | 6 | 0.25 ± 0.40 | 31 |

| | | | | | | | | | | | | | | | | | | | | |
|---|---|---|---|---|---|---|---|---|---|---|---|---|---|---|---|---|---|---|---|---|
| E7 | Cocci | 2.6 | 2.4 | 5.1 | 1.4 ± 0.027 | 2 | 1.4 ± 0.033 | 53 | / | / | / | / | / | / | / | / | 0.46 ± 0.18 | 5 | 0.20 ± 0.17 | 19 |
| E8 | Cocci | 2.2 | 2.2 | 3.7 | 0.93 | 1 | 0.82 ± | 22 | / | / | / | / | / | / | / | / | 0.45 ± 0.27 | 8 | 0.15 ± 0.13 | 32 |
| E9 | Cocci | 2.2 | 1.7 | 3.9 | 1.6 | 1 | 2.0 ± | 52 | 0.40 ± 0.12 | 4 | 0.14 ± 0.065 | 14 | / | / | / | / | 0.37 ± 0.21 | 3 | 0.11 ± 0.11 | 8.7 |
| E10 | Cocci | 2.3 | 2.0 | 3.7 | 0.99 ± 0.042 | 2 | 0.76 ± 0.075 | 42 | / | / | / | / | / | / | / | / | 1.0 ± 0.17 | 2 | 0.66 ± 0.019 | 36 |
| F1 | Rod | 7.2 | 1.9 | 12 | 1.4 ± 0.23 | 2 | 1.3 ± 0.46 | 23 | / | / | / | / | / | / | / | / | 0.73 ± 0.11 | 11 | 0.38 ± 0.14 | 36 |
| F2 | Rod | 4.6 | 1.5 | 5.6 | 1.5 ± 0.75 | 2 | 1.5 ± 1.3 | 55 | 0.61 ± 0.322 | 2 | 0.30 ± 0.257 | 11 | / | / | / | / | 0.66 ± 0.13 | 4 | 0.32 ± 0.15 | 22 |
| F3 | Rod | 5.6 | 1.5 | 8.2 | 1.8 ± 0.14 | 2 | 2.5 ± 0.062 | 62 | / | / | / | / | / | / | / | / | 0.64 ± 0.20 | 10 | 0.32 ± 0.16 | 39 |
| F4 | Rod | 8.6 | 1.9 | 14 | 1.7 ± 0.0040 | 2 | 1.9 ± 0.13 | 27 | / | / | / | / | / | / | / | / | 0.66 ± 0.15 | 10 | 0.38 ± 0.12 | 26 |
| F5 | Rod | 4.7 | 1.0 | 3.6 | 1.2 ± 0.059 | 2 | 0.78 ± 0.036 | 43 | 0.10 ± 0.0090 | 2 | 0.0085 ± 0.0025 | 0.4 7 | / | / | / | / | 0.35 ± 0.059 | 4 | 0.08 ± 0.02 | 9.1 |
| F6 | Rod | 5.7 | 1.9 | 9.4 | 0.47 ± 0.013 | 2 | 0.13 ± 0.026 | 2.8 | / | / | / | / | / | / | / | / | 0.47 ± 0.19 | 13 | 0.20 ± 0.14 | 27 |
| F7 | Rod | 6.6 | 1.8 | 12 | 0.93 ± 0.062 | 2 | 0.58 ± 0.079 | 9.7 | / | / | / | / | / | / | / | / | / | / | / | / |
| F8 | Rod | 4.9 | 2.1 | 9.0 | / | / | / | / | / | / | / | / | / | / | / | / | / | / | / | / |
| F9 | Rod | 5.5 | 2.1 | 8.8 | / | / | / | / | 0.6 | 1 | 0.26 | 2.9 | / | / | / | / | 0.40 ± 0.166 | 5 | 0.13 ± 0.073 | 7.1 |
| F10 | Spirilla | 3.4 | 1.0 | 3.0 | / | / | / | / | 0.10 ± 0.049 | 8 | 0.011 ± 0.011 | 2.9 | / | / | / | / | 0.77 | 1 | 0.44 | 5 |
| F11 | Spirilla | 4.8 | 1.5 | 4.7 | / | / | / | / | / | / | / | / | / | / | / | / | 0.15 ± 0.030 | 16 | 0.018 ± 0.0084 | 9.6 |
| F12 | Cocci | 2.2 | 1.6 | 3.1 | 1.2 ± 0.065 | 2 | 1.2 ± 0.04 | 76 | 0.19 ± 0.049 | 2 | 0.027 ± 0.013 | 1.8 | / | / | / | / | 0.26 ± 0.054 | 6 | 0.060 ± 0.025 | 4.1 |
| F13 | Cocci | 2.0 | 2.1 | 3.4 | 1.2 ± 0.087 | 2 | 0.89 ± 0.20 | 52 | / | / | / | / | / | / | / | / | 0.57 ± 0.016 | 2 | 0.24 ± 0.017 | 16 |

| | | | | | | | | | | | | | | | | | | | | |
|---|---|---|---|---|---|---|---|---|---|---|---|---|---|---|---|---|---|---|---|---|
| F14 | Cocci | 2.8 | 2.3 | 5.5 | 1.3 ± 0.0065 | 2 | 0.67 ± 0.66 | 24 | / | / | / | / | / | / | / | / | 0.92 | 1 | 0.63 | 13 |
| F15 | Cocci | 2.0 | 1.8 | 2.5 | 0.77 ± 0.025 | 2 | 0.43 ± 0.0020 | 35 | / | / | / | / | / | / | / | / | 0.41 ± 0.14 | 2 | 0.14 ± 0.084 | 8.8 |
| F16 | Cocci | 2.4 | 2.3 | 4.6 | 0.9 | 1 | 0.61 ± | 13 | 0.4 | 1 | 0.139 | 3.1 | / | / | / | / | 0.51 ± 0.332 | 5 | 0.29 ± 0.40 | 43 |
| F17 | Cocci | 2.4 | 1.7 | 3.5 | 0.32 | 1 | 0.069 | 2 | / | / | / | / | / | / | / | / | 0.44 ± 0.068 | 4 | 0.14 ± 0.050 | 10 |
| F18 | Cocci | 2.4 | 1.9 | 4.0 | 0.26 ± 0.010 | 2 | 0.048 ± 0.00050 | 2.4 | 0.19 ± 0.044 | 6 | 0.030 ± 0.014 | 4.5 | / | / | / | / | 0.17 ± 0.039 | 6 | 0.03 ± 0.0073 | 6.4 |
| F19 | Cocci | 1.8 | 1.9 | 2.8 | 0.29 ± 0.037 | 2 | 0.071 ± 0.025 | 5.1 | / | / | / | / | / | / | / | / | / | / | / | / |
| F20 | Cocci | 1.7 | 1.6 | 2.3 | / | / | / | / | 0.12 | 1 | 0.009 | 0.4 | / | / | / | / | 0.44 ± 0.0075 | 2 | 0.14 ± 0.015 | 6.2 |
| F21 | Cocci | 1.7 | 1.7 | 2.2 | / | / | / | / | 0.21 | 1 | 0.033 | 1.5 | / | / | / | / | 0.36 | 1 | 0.11 | 3.1 |
| F22 | Cocci | 1.8 | 1.4 | 2.1 | / | / | / | / | 0.18 ± 0.060 | 5 | 0.034 ± 0.009 | 8.1 | / | / | / | / | 0.25 | 1 | 0.04 | 1.1 |
| F23 | Cocci | 1.8 | 1.6 | 2.3 | / | / | / | / | / | / | / | / | / | / | / | / | 0.76 | 1 | 0.51 | 18 |
| F24 | Cocci | 1.7 | 1.6 | 2.5 | / | / | / | / | / | / | / | / | / | / | / | / | / | / | / | / |

[a] MTB ID number is corresponding to the labeling of MTB cells in Fig. 2.

Table S4. Characteristics of the constructed MTB genomes. Genomes were recovered from magnetically enriched MTB cells.

| Parameter | Genome size (bp) | No. of scaffolds | Largest scaffold (bp) | GC (%) | Completeness | Contamination | Quality |
|---|---|---|---|---|---|---|---|
| LS_bin1 | 3996786 | 3822 | 2508 | 39 | 94 | 0.84 | High quality |
| LS_bin2 | 3309862 | 3545 | 2560 | 37 | 66 | 0.84 | Medium quality |
| LS_bin3 | 2778182 | 3571 | 1393 | 37 | 52 | 1.7 | Medium quality |
| LS_bin4 | 2873692 | 3232 | 4338 | 35 | 90 | 2.7 | High quality |
| LS_bin5 | 3961044 | 3801 | 6422 | 35 | 100 | 1.0 | High quality |
| LS_bin6 | 2268743 | 2519 | 1327 | 39 | 62 | 2.0 | Medium quality |
| BS_bin1 | 3508397 | 3285 | 7525 | 38 | 96 | 2.2 | High quality |
| BS_bin2 | 7248057 | 5466 | 10417 | 38 | 97 | 1.1 | High quality |

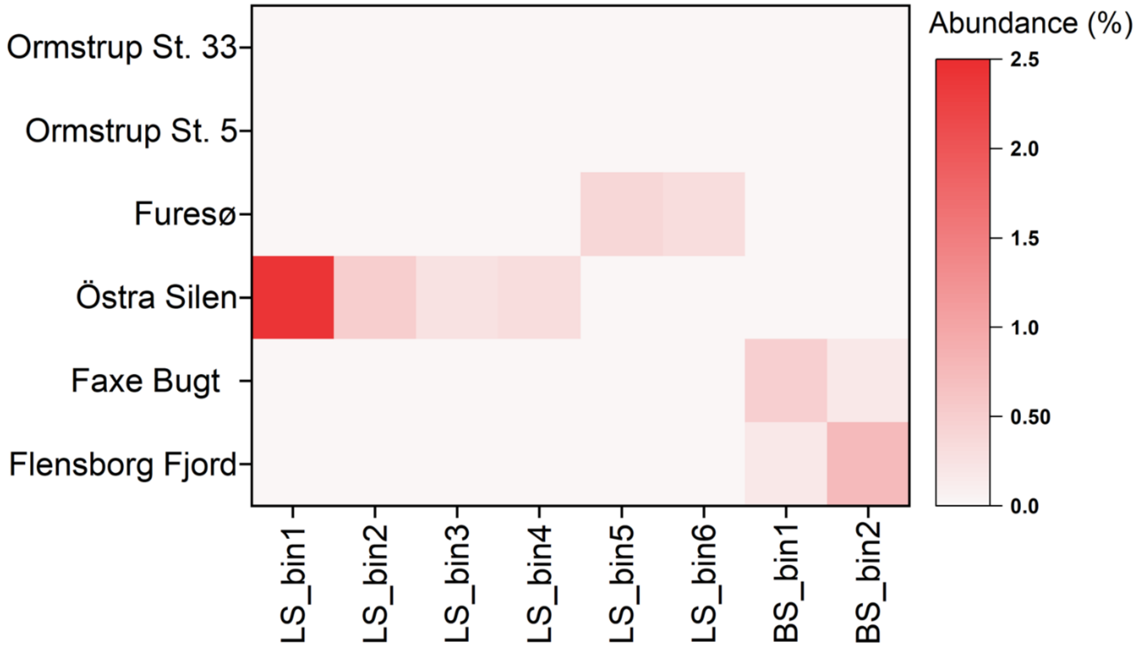


Fig. S5. Relative abundance of the constructed MTB genomes. The relative abundance of genomes in magnetically enriched MTB samples was calculated by normalizing the number of reads mapped to each genome against the total mapped reads in the sample. The abbreviations “LS_bin” and “BS_bin” represent MTB genomes recovered from lake and brackish sediments, respectively.

Table S5. Nomenclature of the constructed MTB genomes.

| MTB genomes | Domain | Phylum | Class | Order | Family | Genus |
|---|---|---|---|---|---|---|
| LS_bin1 | Bacteria | Pseudomonadota | Magnetococcia | / | / | / |
| LS_bin2 | Bacteria | Pseudomonadota | Magnetococcia | / | / | / |
| LS_bin3 | Bacteria | Pseudomonadota | Magnetococcia | / | / | / |
| LS_bin4 | Bacteria | Pseudomonadota | Alphaproteobacteria | Rhodospirillales | *Rhodospirillaceae* | *Magnetospirillum* |
| LS_bin5 | Bacteria | Pseudomonadota | Alphaproteobacteria | Rhodospirillales | *Rhodospirillaceae* | *Magnetospirillum* |
| LS_bin6 | Bacteria | Thermodesulfobacteriota | Desulfobacteria | Desulfobacterales | *Desulfobacteraceae* | *Desulfamplus* |
| BS_bin1 | Bacteria | Pseudomonadota | Magnetococcia | / | / | / |
| BS_bin2 | Bacteria | Thermodesulfobacteriota | Desulfobacteria | Desulfobacterales | *Candidatus* Magnetomoraceae | *Candidatus* Magnetomorum |

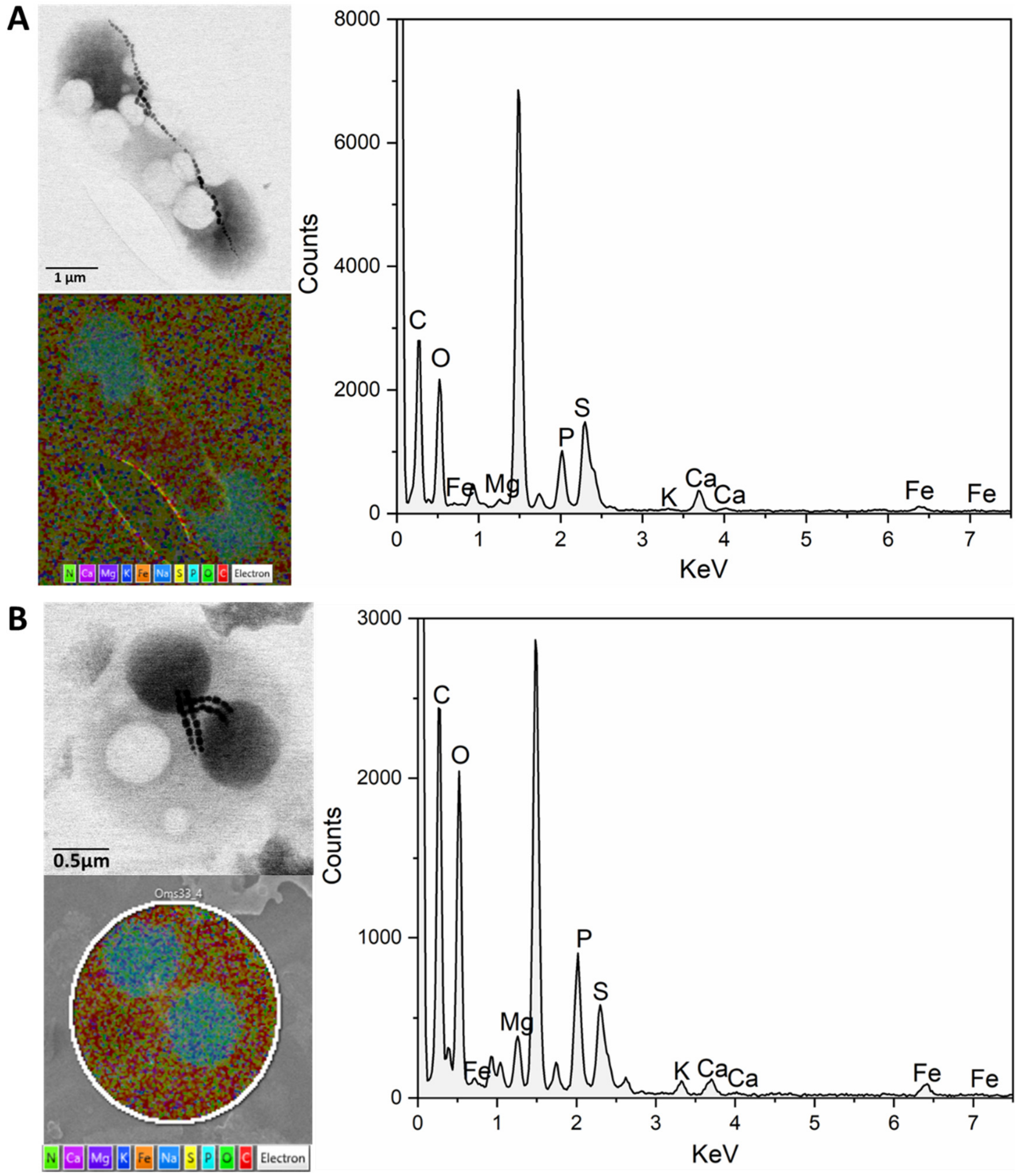


Fig. S6. Examples of STEM-EDX mapping and spectrum illustration of intracellular (A) Ca-rich polyP inclusions and (B) Mg-rich polyP inclusions within MTB cells. Cells were magnetically enriched from Ormstrup St. 33 sediment.

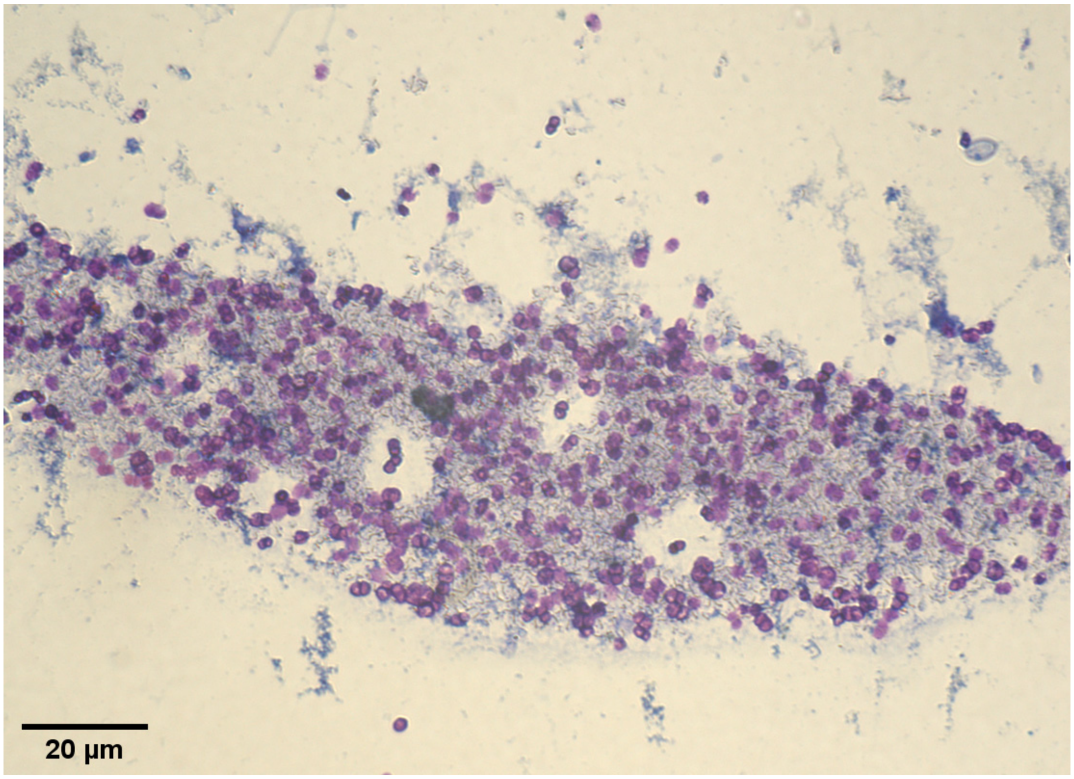


Fig. S7. Example of toluidine staining of magnetically collected MTB cells from Faxe Bugt sediment. The dark red-purple color was observed as a result of a metachromatic effect occurring when toluidine blue binds to polyphosphate.

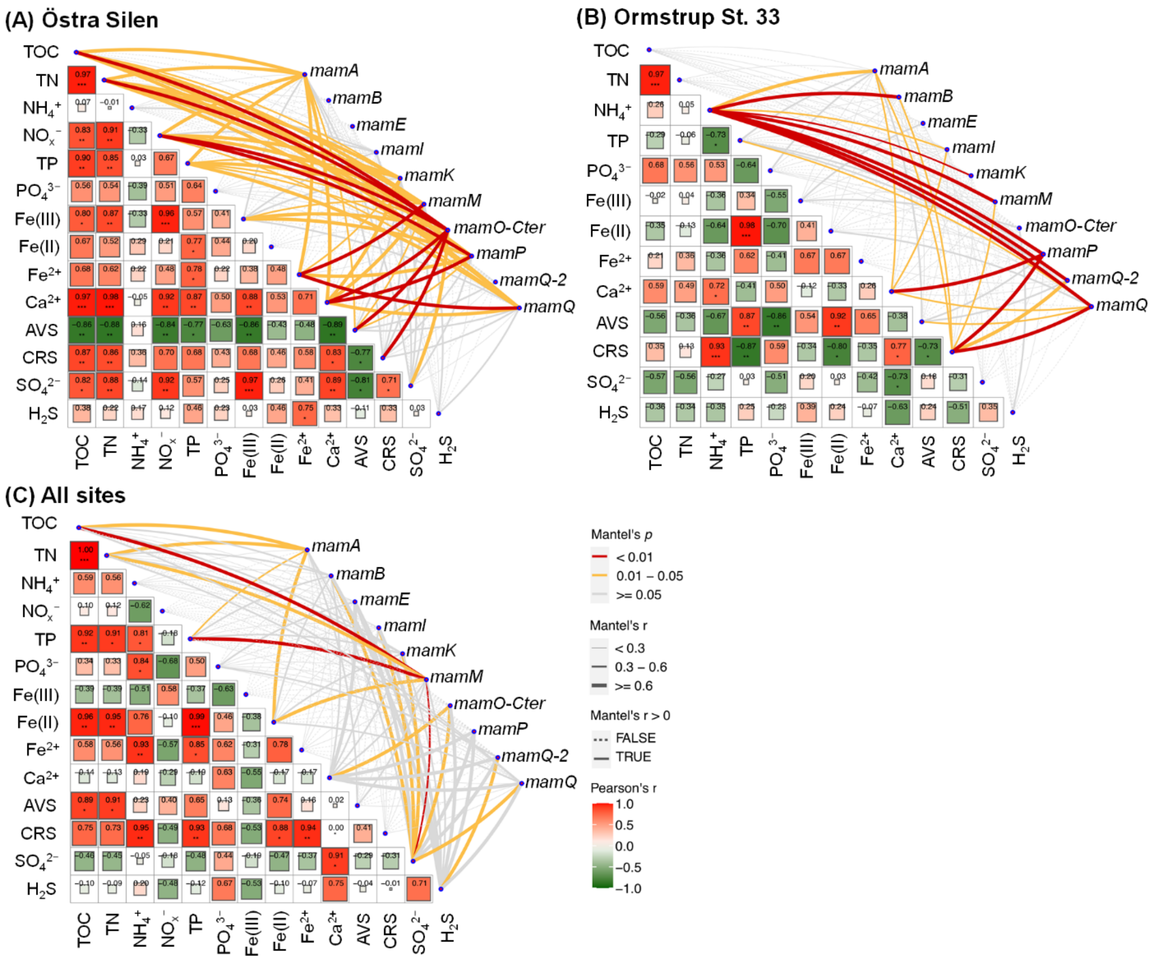


Fig. S8. Correlations of the depth distribution of *mam* genes with geochemical parameters: (A) Östra Silen, (B) Lake Ormstrup St. 33, and (C) All sampling sites. The red and yellow lines represent significant correlations of *mam* genes with geochemical parameters (Montel's test, $p < 0.05$), while grey line indicate insignificant correlations (Montel's test, $p > 0.05$). The thicker line indicates stronger correlation. The red and green squares represent positive and negative correlations (Pearson's test) between geochemical parameters, respectively. Bigger square size and higher correlation coefficient value indicate stronger correlation, and vice versa. Correlation coefficient values marked with asterisk (*) and (**) represent significant correlations with $p$ values < 0.05 and <0.01, respectively. The depth distribution of geochemical parameters and *mam* genes in sediments is shown in Fig. S2 and Fig. 1C, respectively.

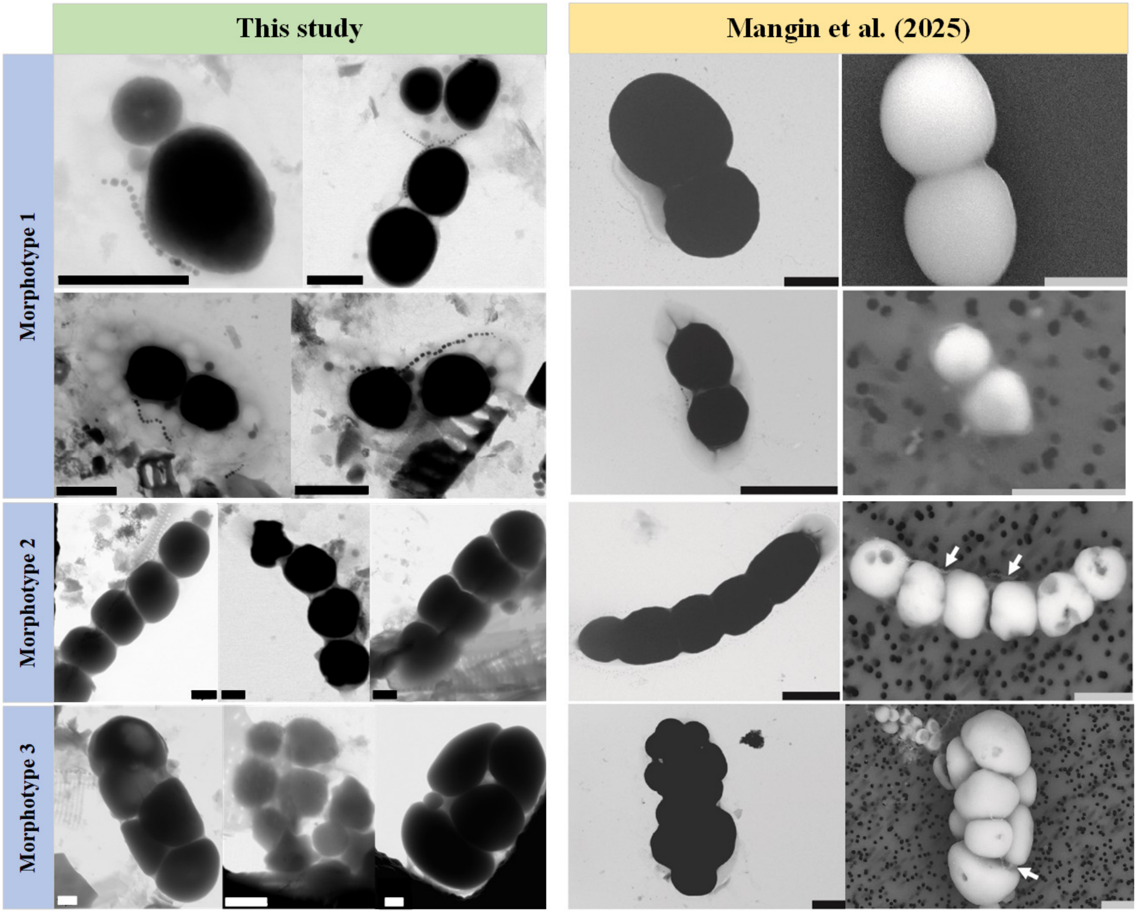


Fig. S9. Comparison of morphotypes of $CaCO_3$ inclusions in MTB identified in our study and Mangin et al. (2025).

Table S6. Estimation of storage and annual yield of polyP inclusions, S globules, and $NO_3^-$ vacuoles in MTB cells in the investigated freshwater sediments.

| **Parameter** | **Value** |
| --- | --- |
| Sediment depth existing MTB (cm) | 8 |
| Average number of MTB cells per mL | $10^3$–$10^6$ |
| Generation turnover time of environmental MTB (h) | 12 |
| Average MTB generations per year | 730 |
| **polyP inclusion** [a] | |
| Percentage of MTB with polyP inclusions | 0.75 |
| Average number of polyP inclusions per MTB cell | 3.85 |
| Average diameter of polyP inclusions (μm) | 1.06 |
| Mass of single polyP inclusion in 1.06 μm diameter (g P) [b] | $3.93\times10^{-14}$ |
| Total mass of polyP stored in MTB (kg P/cm$^2$) | $1.82\times(10^{-12}$–$10^{-9})$ |
| Annual yield of polyP inclusions by MTB (kg P/cm$^2$) | $6.63\times(10^{-10}$–$10^{-7})$ |
| **$NO_3^-$ vacuoles** [a] | |
| Percentage of MTB with $NO_3^-$ vacuoles | 0.67 |
| Average number of $NO_3^-$ vacuoles per MTB cell | 6.14 |
| Average diameter of $NO_3^-$ vacuoles (μm) | 0.46 |
| Mass of single $NO_3^-$ vacuole in diameter of 0.46 μm (g N) [b] | $7.33\times10^{-14}$ |
| Total N mass stored in MTB (kg N/cm$^2$) | $4.83\times(10^{-12}$–$10^{-9})$ |
| Annual yield of $NO_3^-$ vacuoles (kg N/cm$^2$) | $1.76\times(10^{-9}$–$10^{-3})$ |
| **S globules** [a] | |
| Percentage of MTB with S globules | 0.25 |
| Average number of S globules per MTB cell | 5.06 |
| Average diameter of S globules (μm) | 0.28 |
| Mass of single S globule in diameter of 0.28 μm (g S) [b] | $1.38\times10^{-14}$ |
| Total S mass stored in MTB (kg S/cm$^2$) | $7.28\times(10^{-13}$–$10^{-10})$ |
| Annual yield of S globules (kg S/cm$^2$) | $2.66\times(10^{-10}$–$10^{-7})$ |

[a] Annual production of intracellular inclusions by MTB is estimated using the formula: (sediment depth existing MTB) × (average number of MTB cells per mL) × (percentage of MTB with inclusions) × (average number of inclusions per MTB cell) × (mass of single inclusions) × (average number of MTB generations per year). [b] Values were adapted from Lin et al. (2014), Schulz-Vogt et al. (2019), Dahl (2020), Schulz and Jørgensen (2001), etc.

Table S7. Proportion of intracellular stored P, N, and S in MTB to P, N, and S pools in the investigated freshwater sediments.

| **P, N, and S pools in investigated freshwater sediments [a]** | | | |
|---|---|---|---|
| Sampling site | TP (kg/cm$^2$) | TN (kg/cm$^2$) | TS (kg/cm$^2$) |
| Östra Silen | $4.68\times10^{-7}$ | $6.95\times10^{-7}$ | $3.17\times10^{-7}$ |
| Furesø | $3.03\times10^{-7}$ | $5.30\times10^{-7}$ | $1.33\times10^{-6}$ |
| Ormstrup St. 5 | $1.22\times10^{-6}$ | $1.22\times10^{-5}$ | $4.89\times10^{-6}$ |
| Ormstrup St. 33 | $1.83\times10^{-6}$ | $1.02\times10^{-5}$ | $1.00\times10^{-5}$ |
| **Intracellular stored P, N, and S by MTB [b]** | | | |
| Mass stored in MTB | polyP (kg P/cm$^2$) | $NO_3^-$ (kg N/cm$^2$) | S (kg S/cm$^2$) |
| | $1.82\times(10^{-12}–10^{-9})$ | $4.83\times(10^{-12}–10^{-9})$ | $7.28\times(10^{-13}–10^{-10})$ |
| **Proportion of intracellular stored P, N, and S in MTB cells to sediment P, N, and S pools** | | | |
| Sampling site | P (%) | N (%) | S (%) |
| Östra Silen | $3.88\times(10^{-4}–10^{-1})$ | $6.95\times(10^{-4}–10^{-1})$ | $2.30\times(10^{-4}–10^{-1})$ |
| Furesø | $6.00\times(10^{-4}–10^{-1})$ | $9.11\times(10^{-4}–10^{-1})$ | $5.45\times(10^{-5}–10^{-2})$ |
| Ormstrup St. 5 | $1.49\times(10^{-4}–10^{-1})$ | $3.97\times(10^{-5}–10^{-2})$ | $1.49\times(10^{-5}–10^{-2})$ |
| Ormstrup St. 33 | $9.92\times(10^{-5}–10^{-2})$ | $4.73\times(10^{-5}–10^{-2})$ | $7.27\times(10^{-6}–10^{-3})$ |

[a] Values were integrated over 0 to 8 cm depth stratum. Total S is the sum of integrated values of AVS, CRS, and $H_2S$. [b] The detail of the calculations refers to Table S6.

### Supplementary Note-Analyses of sediment solid-phase and porewater

#### Benthic oxygen uptake rate

Four sediment cores from each site were submerged carefully in a continuous aerated tank with on-site water, and pre-incubated for >24 hours in the dark at near *in situ* temperature. After pre-incubation, the height of the water column above the sediment was measured and the sediment cores were sealed with rubber stoppers, leaving no air bubbles. Dissolved $O_2$ concentrations were measured every half hour with PyroScience noninvasive optical oxygen sensors (Germany) until $O_2$ concentrations reached ~50% saturation. The total benthic $O_2$ uptake rate as a proxy for total carbon mineralization rate (mmol/m$^2$/d) (Canfield et al., 1993a) was calculated as the linear regression slope of $O_2$ concentration, accounting for the enclosed area and water volume.

#### Sediment characteristics and solid-phase analysis

Porosity was calculated by multiplying wet density (weight of a known volume) and water content (weight loss after drying the sediment samples at 105 °C for 24 hours). The grain-size distribution of sediments was determined using a Hydro Mastersizer 3000 (Malvern). Each sample was run in three times to check for floc destruction and possible variance. Sub-samples for solid-phase Fe(III) and Fe(II) concentrations were added to 15 ml tubes leaving little headspace and stored frozen under $N_2$ atmosphere until cold HCl extraction (Kostka and Luther, 1994). This assay extracts poorly crystalline Fe(III) oxides and particulate Fe(II), such as FeS and $FeCO_3$ (Canfield et al., 1993b; Thamdrup et al., 1994). The extractions were performed with 10 ml extractant and ~50−150 mg wet sediment. Extraction time was 1 h in the dark. The concentrations of the oxidation stages of Fe were determined separately by using a Ferrozine solution (50 mmol/L HEPES, 0.1% Ferrozine, pH 7), with and without 1% (w/v) hydroxylamine hydrochloride for the quantification of total Fe and Fe(II), respectively (Canfield et al., 1993b). Homogenized sediments were fixed in 20% zinc acetate (vol:vol = 1:1) and stored frozen until analysis. The two-step distillation method was used to separate AVS (iron sulfide (FeS) + $H_2S$) from CRS ($S^0$ + iron disulphide ($FeS_2$)) (Troelsen and Jørgensen, 1982). Sulfide concentrations were determined spectrophotometrically by the methylene blue technique (Cline, 1969). Samples for TP, TOC and TN was stored frozen without pre-treatment until analysis. To determine the TP content, ~0.3 g of sediment was dried for 24 hours at 105 °C followed by extraction in 10 ml 65% $HNO_3$ in a high-pressure microwave (400 W, 800 PSI, 200 °C, 15 min ramp, and 15 min hold time; CEM with MarsXpress vessels). The concentration of TP was measured as dissolved phosphate ($PO_4^{3-}$) in the extract using Inductively Coupled Plasma with Optical Emission Spectroscopy (ICP-OES; Optima 2100 DV, Perkin Elmer, Shelton, Connecticut, USA). Before the analysis of TOC and TN, the sediment was dried at 105 °C for 24 hours and acidified with 1 M HCl to remove inorganic carbon. Subsequently, 15–25 mg of dried sediment was packed in tin capsules and combusted on an elemental analyzer (Flash 2000, Thermo Fisher Scientific).

### Preservation and measurement of porewater samples

Samples for the analysis of $H_2S$ and $SO_4^{2-}$ were collected in 2 ml vials, preserved with 20 µL of 0.5% or 20% (w/v) zinc acetate depending on salinity, and frozen for later analysis. Aliquots of 2 mL were acidified separately: 20 µL of 6 M HCl was added for $Fe^{2+}$ determination, 20 µL of 2 M $H_2SO_4$ was added for ortho-$PO_4^{3-}$ analysis, and 20 µL of 65% $HNO_3$ was added for dissolved $Ca^{2+}$ analysis. The rest of the filtered porewater was immediately frozen to quantify dissolved inorganic N species ($NH_4^+$, $NO_2^-$, and $NO_3^-$).

Porewater $H_2S$ concentrations were measured colorimetrically using the methylene blue method (Cline, 1969), while $SO_4^{2-}$ concentrations were quantified by suppressed anion chromatography (Dionex). The concentration of $Fe^{2+}$ was quantified colorimetrically using Ferrozine without reducing agent (Stookey, 1970; Thamdrup et al., 1994). Ortho-$PO_4^{3-}$ concentrations were determined spectrophotometrically by the molybdenum-reactive P method

(Murphy and Riley, 1962). Dissolved $Ca^{2+}$ concentrations were measured by ICP-OES (7700x, Agilent Technologies). An air-segmented continuous-flow analyzer (SKALAR San$^{++}$, Netherlands) was used for colorimetric analysis of $NH_4^+$, $NO_2^-$, and $NO_x^-$ (i.e., $NO_2^-$ + $NO_3^-$) concentrations.
